# Uncovering missed indels by leveraging unmapped reads

**DOI:** 10.1101/488601

**Authors:** Mohammad Shabbir Hasan, Xiaowei Wu, Liqing Zhang

## Abstract

In current practice, Next Generation Sequencing (NGS) applications start with mapping/aligning short reads to the reference genome, with the aim of identifying genetic mutations. While most short reads can be mapped to the reference genome accurately by existing alignment tools, a significant number remain unmapped and excluded from downstream analyses thus potentially discarding important biological information hidden in the unmapped reads. This paper describes Genesis-indel, a computational pipeline that explores the unmapped reads to identify novel indels that are initially missed in the alignment procedure. Genesis-indel is applied to the unmapped reads of 30 Breast Cancer patients from TCGA. Results show that the unmapped reads are conserved between the two subtypes of breast cancer investigated in this study and might contribute to the divergence between the subtypes. Genesis-indel is able to leverage the unmapped reads to identify 72,997 small to large novel high-quality indels previously not found in the original alignments and among them, 16,141 have not been annotated in the widely used mutation database. Statistical analysis shows that these new indels mostly altered the oncogenes and tumor suppressor genes. Functional annotation further reveals that these indels are strongly correlated to pathways of cancer and can have high to moderate impact on protein functions. Additionally, these indels overlap with the genes that are missed in the indels from the originally mapped reads and contribute to the tumorigenesis in multiple carcinomas.

## Introduction

Next Generation Sequencing (NGS) facilitates generation of an enormous number of short reads and allows the identification of genomic mutations that cause phenotype changes and genetic diseases such as Mendelian disorders [1], Acute Myeloid Leukemia [2], and Lung cancer [3]. Applications analyzing the NGS reads typically start with mapping the short reads against a reference genome and then based on the mapped reads, determine the genetic mutations such as Single Nucleotide Polymorphism (SNP) and sequence variants such as Insertion and Deletion (indel) of bases. Many alignment algorithms have been developed to map the short reads to the reference genome such as MAQ [4], SOAP [5], BWA [6], Bowtie [7], Bowtie2 [8], SNAP [9], and SOAP2 [10], to name a few. Although these alignment tools are very efficient in aligning the short reads, a nonnegligible fraction of reads are left unmapped due to (1) structural variants longer than the allowed number of gaps and mismatches by the mapper, (2) sequencing error, or (3) sample contamination [11]. In current practice, these unmapped reads are not used for variant calling and downstream analyses, and thus mutations harbored in these unmapped reads remain hidden from any inference on important genotype and phenotype and/or their associations with any disease such as cancer. However, as shown in Figure 1, some of the “hidden” or “missing” mutations can contain the key for understanding the molecular mechanisms of genetic diseases or cancer and might be used as markers for disease/cancer diagnosis and prognosis.

**Figure 1:**
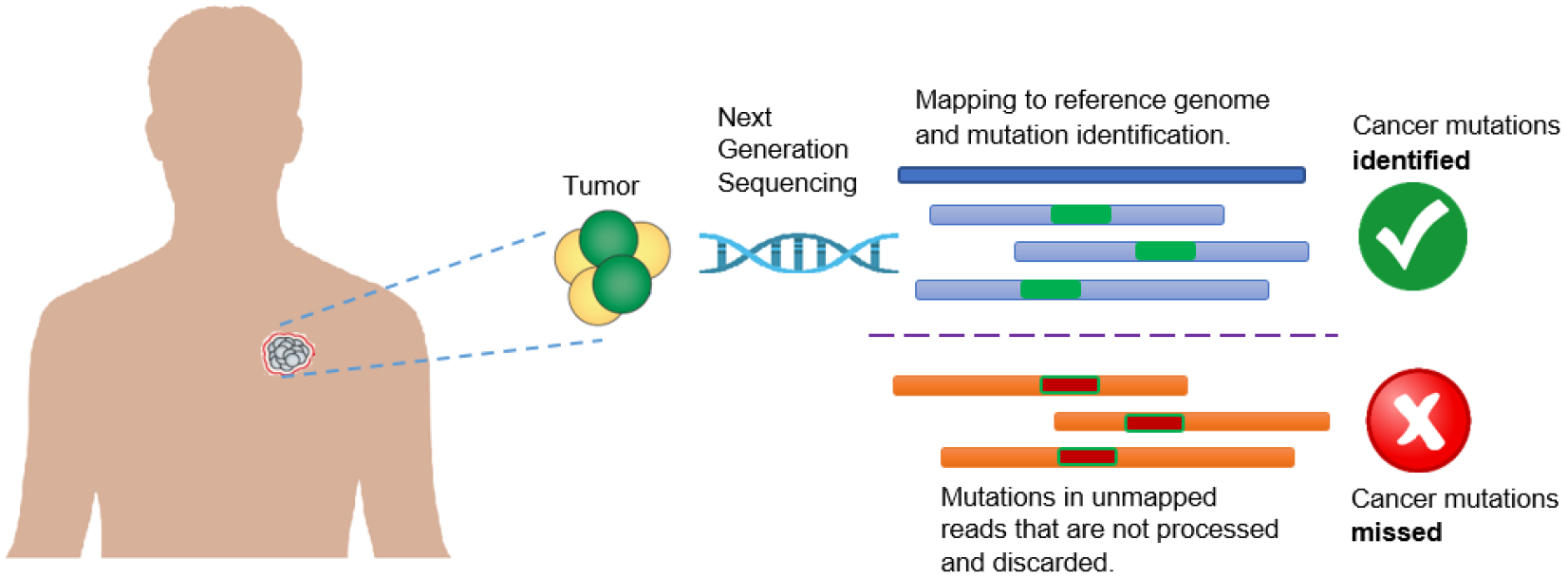
Limitation of current practice in cancer research which discards unmapped reads and therefore misses important mutations containing real biological signal.

The consequence of missing the mutations contained in the unmapped reads can lead to inaccurate downstream analyses such as characterizing the tumor evolution in a cancer patient. Some of these missed mutations can be the hallmark of tumors and can be useful for targeted therapy. Therefore, it is critical to identify the mutations in those regions for clinical decision-making as well as for guided personal treatment [12, 13]. With this objective in mind, it is essential to inspect the unmapped reads previously excluded from analyses to ensure that none of these essential mutations are missed in those regions of interest.

This paper describes Genesis-indel, a computational pipeline to explore unmapped reads for the systematic identification of indels missed in the original alignments. Note that this pipeline focuses on indels only, the second most abundant form of genetic variation in human populations [14–16]. Despite being a common form of genetic variation in humans, indels have not been studied as thoroughly as SNPs, though they have been identified playing a key role in causing diseases such as Cystic fibrosis [17], Fragile X Syndrome [18], acute myeloid leukemia [2, 19, 20], and lung cancer [21]. In addition, insertion of transposable elements such as Alu can affect gene function and change gene expression [22]. Genesis-indel is applied to explore unmapped reads of 30 breast cancer patients from The Cancer Genome Atlas (TCGA) [23] and identify indels hidden in the unmapped reads of these patient genomes. Results show that unmapped reads can be used to cluster samples to different cancer subtypes. In addition, Genesis-indel can successfully curate the unmapped reads and detect small to large novel high-quality indels that are missed previously and some of these indels are specific to a particular subtype of breast cancer. Functional annotation of the newly identified indels shows that the indels found from unmapped reads are strongly correlated with cancer pathways and may play an important role in cancer progression. In addition, these indels overlap with genes where no mutation was found from the original alignments and these genes can contribute to the tumorigenesis in multiple cancers. Therefore, this study shows great promise in complementing the current procedure of understanding the underlying mechanism of cancer progression and will be useful for clinical decision making.

**Figure 2:**
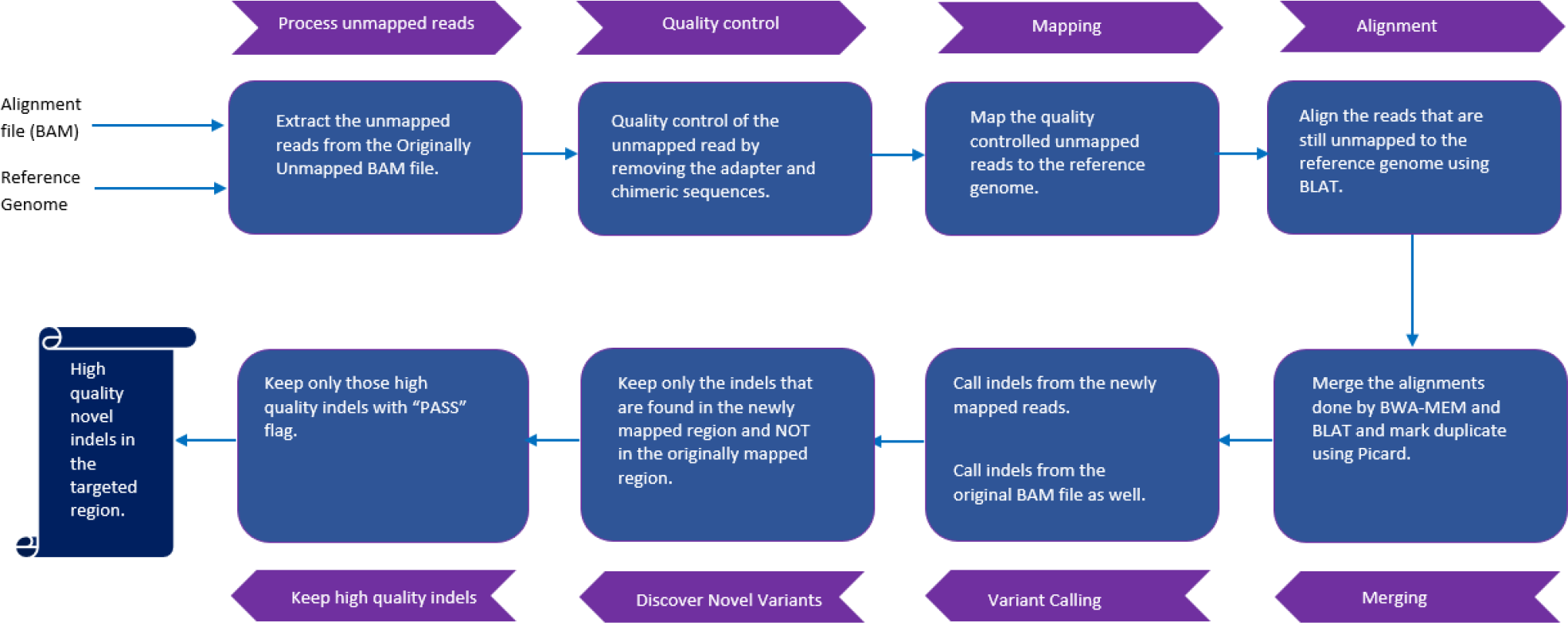
Genesis-indel workflow. The input to Genesis-indel is the alignment file (BAM file) and the reference genome (FASTA format). First, the unmapped reads are extracted from the input BAM and passed to the quality control module. After quality control, the reads are mapped to the reference genome using BWA-MEM. The reads that still remained unmapped are aligned using BLAT. In the merging step, the output of BWA-MEM and BLAT are merged and duplicates are marked using Picard. The merged alignment is then passed to the Variant Calling module followed by quality filtering of the indels. Finally, the output contains novel high-quality indels rescued from the originally unmapped reads.

## Results and Discussion

Figure 2 shows the schematic of the Genesis-indel workflow (see Methods for detail). Genesis-indel is used to identify the novel high-quality indels from the alignment (BAM files) of 30 Breast Cancer patients deposited in TCGA. These BAM files were originally produced by mapping the raw sequencing reads of these patients to the human reference genome using BWA [6].

### Existence of a nonnegligible number of originally unmapped reads

The alignment file of each patient sample is processed by using SAMtools [24] to extract the “Originally Unmapped” reads. For a given individual investigated here, 6.6 to 74 million reads are unmapped (average = 31.86 million). Shown in Figure 3, the unmapped reads constitute an average of 5% of the total reads (altogether there are more than 955 million reads unmapped for 30 patient samples) in the original alignment files provided by TCGA. Genesis-indel targets these discarded reads to rescue the indels missed in the original alignment.

**Figure 3:**
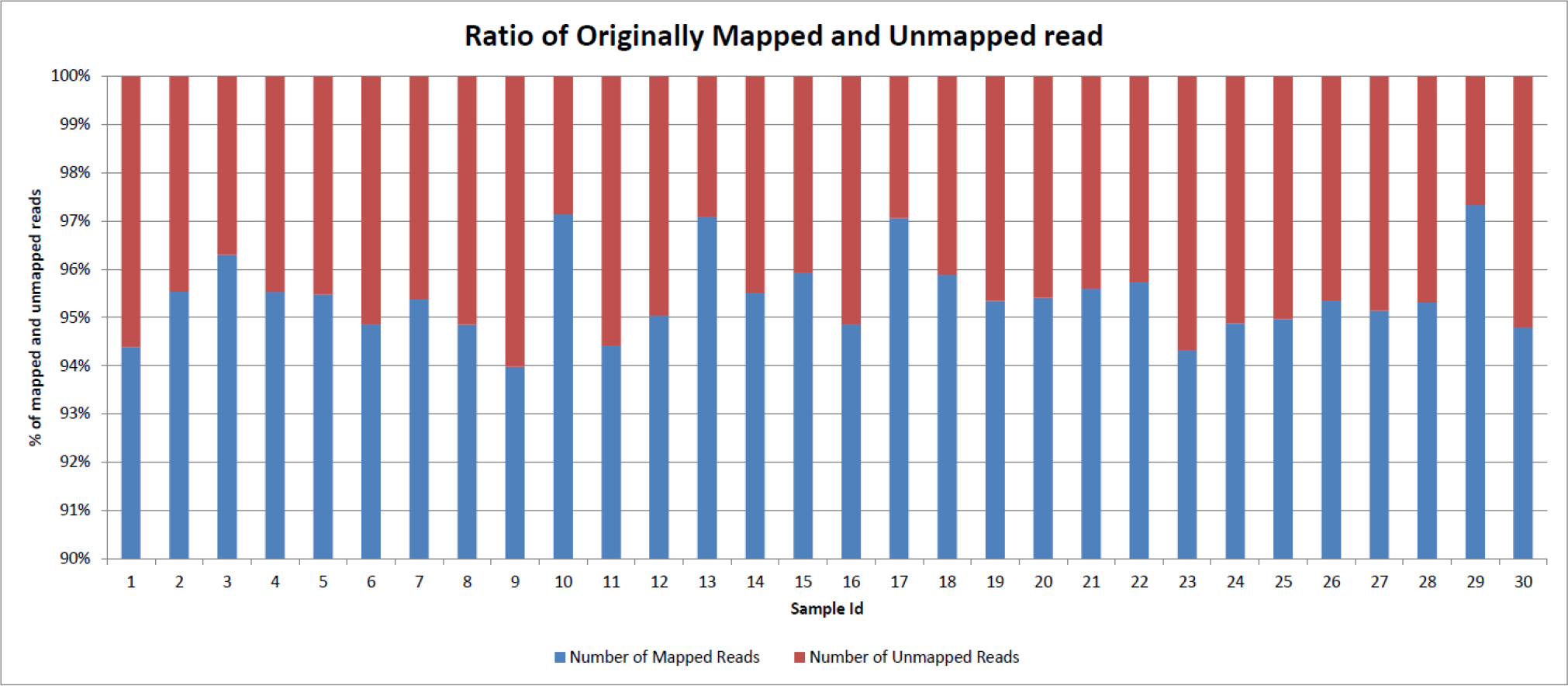
Percentage of mapped and unmapped reads in the original alignment files of the 30 breast cancer patients collected from TCGA.

### Quality control of the unmapped reads

Since it is likely that the unmapped reads might be of low quality and hence ignored by the mapper, the extracted unmapped reads are processed for quality control. The unmapped reads from all samples are combined and passed to FastQC [25] to get various statistics of the reads. According to the report produced by FastQC, the originally unmapped reads have some quality issues such as (1) overall poor per base sequence quality, (2) a poor score for per sequence quality, (3) overrepresentation of “N” contents, (4) overrepresentation of Illumina Paired-End PCR Primer 2 due to PCR over-amplification, and (5) other adapter contents (Supplementary Figures 1(a), 2(a), and 3(a)). In most cases, reads that are contaminated with adapter sequences are simply not mapped because of sequencing errors in the adapter sequences. Therefore, removing these contaminated sequences is expected to improve the quality of the unmapped reads. Trimmomatic [26] is applied to the combined unmapped reads from all samples and then FastQC is used again to assess the quality of the reads. As shown in Supplementary Figures 1(b), 2(b), and 3(b), after trimming adapter sequences, many issues were fixed and the quality of the unmapped reads improved significantly. Although there is a low-quality issue with some k-mer noise at the 3’ end of the reads (Supplementary Figure 4), the mapping is not affected by these k-mers as they are not mapped to the reference genome and hence get discarded during the alignment step. Alternatively, RAUR (Re-align the Unmapped Reads) [27] could be used for read quality control. RAUR analyzes the base quality distribution of sequencing error and adopts an iterative trimming algorithm to remove a segment of low-quality bases from the unmapped read so that the remaining segment of the originally unmapped read can be confidently mapped. After combining RAUR with alignment tools such as BWA and Bowtie2, the alignment rate is improved as observed by Peng et al. [27]. However, RAUR is not publicly available and hence could not be considered for Genesis-indel pipeline. After the quality control by Trimmomatic, for the individuals investigated here, 29.29% to 89.5% of the originally unmapped reads are retained (average = 67.68%) constituting around 647 million reads.

### Mapping the quality controlled unmapped reads

After quality control, the unmapped reads are mapped to the reference genome using BWA-MEM [28]. BWA-MEM can automatically choose between local and end-to-end alignments. It is applicable to map short as well as long reads, and is sensitive in mapping reads with indels. While mapping, unlike other short-read mappers, it allows big gaps potentially caused by structural variants and shows better or comparable performance than several state-of-the-art read mappers to date in terms of speed and accuracy [28]. This mapper is robust to sequencing errors as well. After the reads are aligned by BWA-MEM, some reads still remain unmapped. At this step, another local alignment tool, BLAT (BLAST-Like Alignment Tool) [29] is used to align these reads. After this step, the alignments from BWA-MEM and BLAT are merged. Counting the number of newly mapped reads reveal that 65.38% of the originally unmapped reads (624,892,089 out of 955,822,913) now get mapped to the reference genome. Out of these newly mapped reads, BWA-MEM mapped 479,064,451 reads and BLAT aligned 145,827,638 reads. As mentioned before, the mapper used by TCGA is BWA and although theoretically, BWA can map arbitrarily long reads, practically, the performance in mapping long reads degraded with the increase of the sequencing error rate [27]. By removing the bases with sequencing error and using BWA-MEM which is a robust and variant sensitive mapper coupled with another sensitive aligner BLAT, Genesis-indel manages to map many of the initially unmapped reads. Figure 4 shows the average mapping quality of the newly mapped reads for all samples. For most of the samples, the mapping quality is higher than that of the originally mapped reads (Supplementary Figure 5).

**Figure 4:**
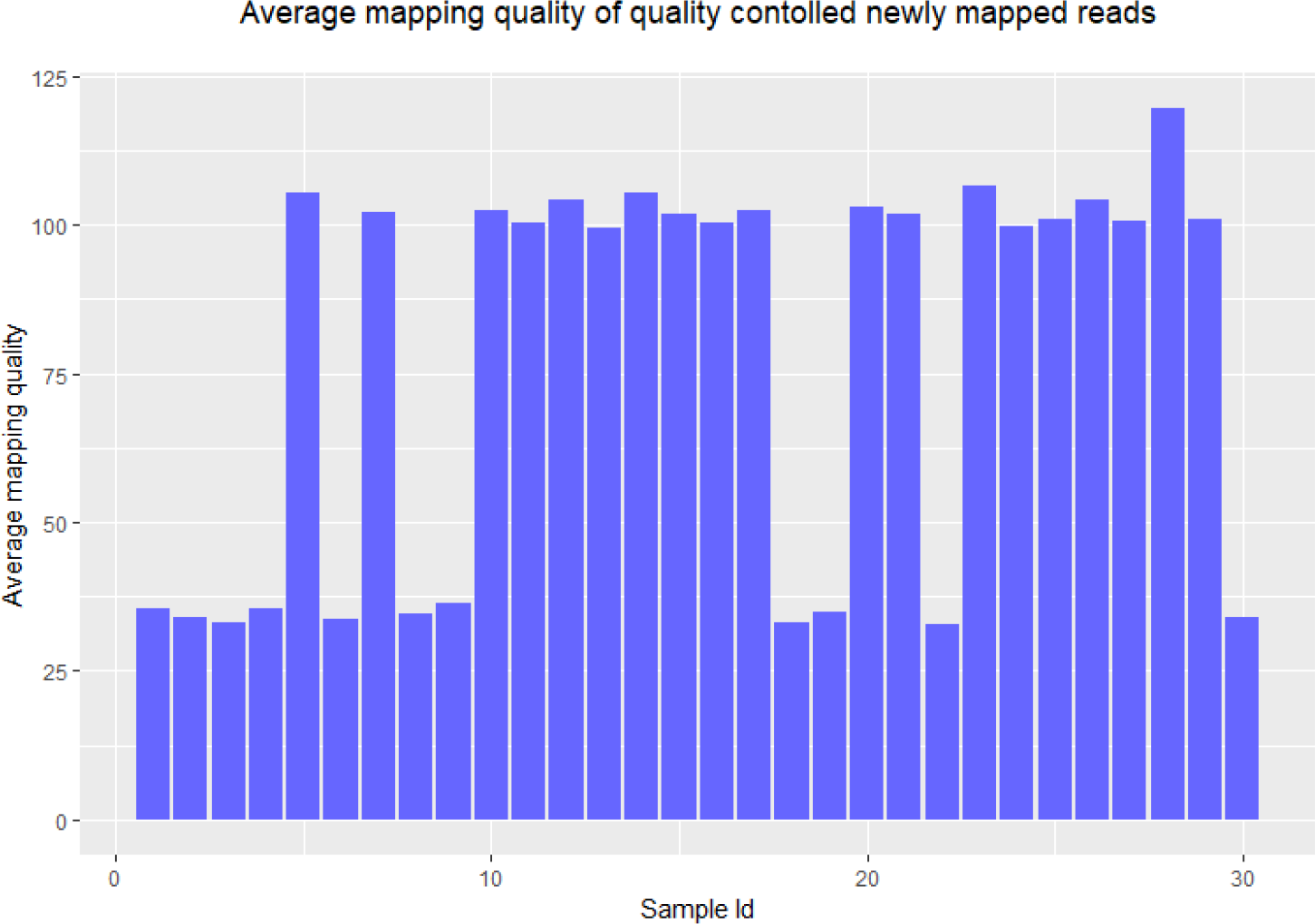
Average mapping quality of the newly mapped reads of all samples.

### Identifying the novel high-quality indels from the newly mapped reads

Genesis-indel uses Platypus [30] to call indels from the newly mapped reads. Separately, indels are also called from the reads that are originally mapped. Platypus is chosen as it performed the best among other existing indel callers based on real data as reported in a recent review [31]. After variant calling, indels from the newly mapped reads are inspected for any match with the indels found in the originally mapped reads. As shown in Figure 5, an indel already in the originally mapped reads can be called again in the newly mapped reads. Since the main objective is to identify novel indels from the originally unmapped reads, these re-identified indels (3.58% of the total number of indels identified from the originally mapped reads) are discarded from the downstream analysis. This step produces 105,331 novel insertions and 143,386 novel deletions from the newly mapped reads. These indels are referred to as “Novel indels” in this paper.

**Figure 5:**
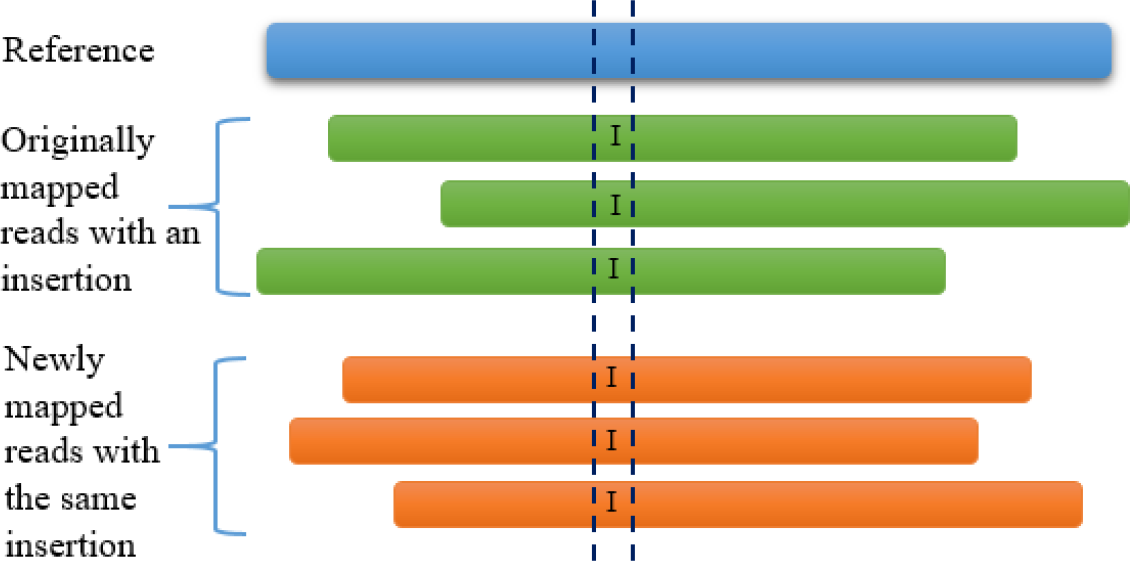
An example of an insertion that is already in the originally mapped reads can be called again in the newly mapped reads. Since this indel can be found in the originally mapped reads and hence not novel by definition, this indel is discarded in the downstream analysis.

Examination of the flags of the novel indels shows that for many of the indels, Platypus does not produce a high confidence value. Therefore, to consider only the high-quality indels for further analysis, novel indels are filtered again to make sure that only high-quality indels are reported finally. Therefore, only novel indels with “PASS” flags are considered for the final result and are termed as “Novel high-quality indel” in this paper. These novel high-quality indels are the ones for which the indel caller has high confidence. In total, Genesis-indel reports 31,924 novel high-quality insertions (43.73% of the total novel high-quality indels) and 41,073 novel high-quality deletions (56.27% of the total novel high-quality indels) from the 30 samples investigated here. The deletion to insertion ratio for the novel high-quality indel is 1.29:1, close to the deletion to insertion ratio for originally mapped reads which is 1.11: 1 (7,313,641 insertions and 8,082,055 deletions).

Figure 6 shows a novel 15 bases deletion in Chromosome 1 that is identified in the newly mapped reads (Figure 6, lower panel) but missed in the original alignment (Figure 6, upper panel). Although this paper focuses on indels only, as shown in Figure 6, in addition to indels, new SNPs are also identified by Genesis-indel in the originally unmapped reads.

**Figure 6:**
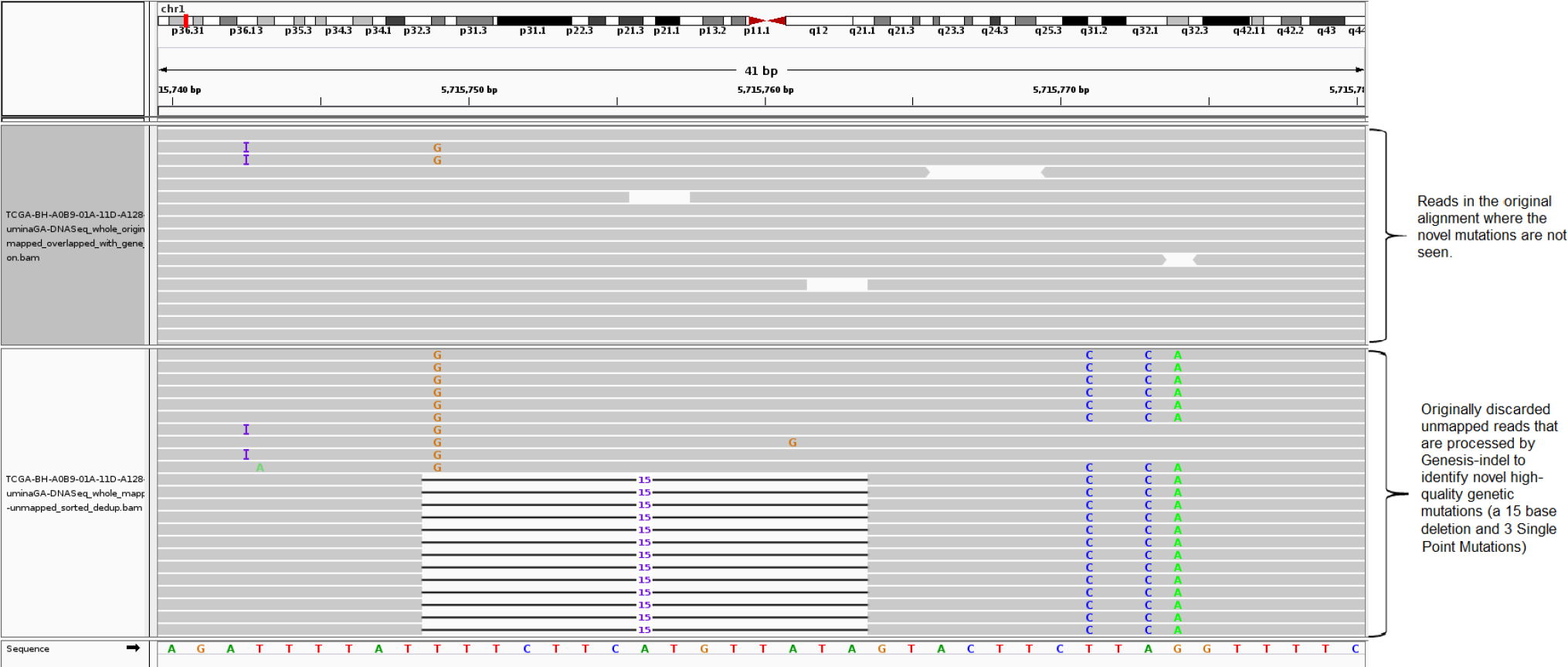
IGV [32] snapshot of a novel high-quality indel identified in the newly mapped read but missed in the original alignment. The upper panel shows the originally mapped reads and the lower shows the newly mapped reads.

### Frameshift indels are more frequent than in-frame types in the novel high-quality indels

Novel high-quality indels identified here contain 53,623 frameshift and 19,374 in-frame indels, indicating a higher abundance of the frameshift indels in the unmapped reads. Frameshift indels are also found more abundant than in-frame indels in the originally mapped reads (13,911,266 vs 1,481,550). Frameshift mutations are common in cancer patients and increase the susceptibility to cancers and other diseases by causing loss of significant fractions of proteins [33–36] which indicates the significance of these newly identified frameshift indels. Frameshift indels of longer length (≥15 bases) are more frequent than in-frame indels of corresponding length (585 insertions, 1,479 deletions vs. 404 insertions, 812 deletions).

Mills et al. studied the landscape of small indels in the genomes of 79 healthy humans and found that 53.5 and 46.5% of these indels are frameshift and in-frame indels, respectively [37]. However, in a study by Iengar et al. [36] on COSMIC indels, it is found that 75.7% and 24.3% of the indels are frameshift and in-frame indels, respectively, which means unlike the distribution of coding indels in the genome of healthy people, frameshift indels are preferred over in-frame indels in cancer genome. Therefore, in addition to the frameshift indels from the originally mapped reads, the novel high-quality frameshift indels uncovered by Genesis-indel may harbor important signals for further analysis.

### Novel high-quality indels have more diverse length than indels in the originally mapped reads

Figure 7 shows the distribution of the length of the novel high-quality indels analyzed here. It is observed that insertion and deletion frequency decreases with the increase in the size of the insertion and deletion. The longest novel high-quality insertion and deletion are 28 and 45 bases, respectively (34 and 55 bases for the indels from the originally mapped reads). It is expected that the novel indels would be long as they might have been missed because the lengths might exceed the number of gaps and mismatches allowed by the mapper. Surprisingly, as shown in Figure 7, most of the newly discovered indels (91% of insertion and 88.1% of deletion) are short indels (≤10 bases) indicating the limitation of the mapper used in the TCGA project. A paired two-tailed t-test shows that the indel length distribution of novel high-quality indels is significantly different from that of indels identified from the originally mapped reads (p-value = 0.00025).

**Figure 7:**
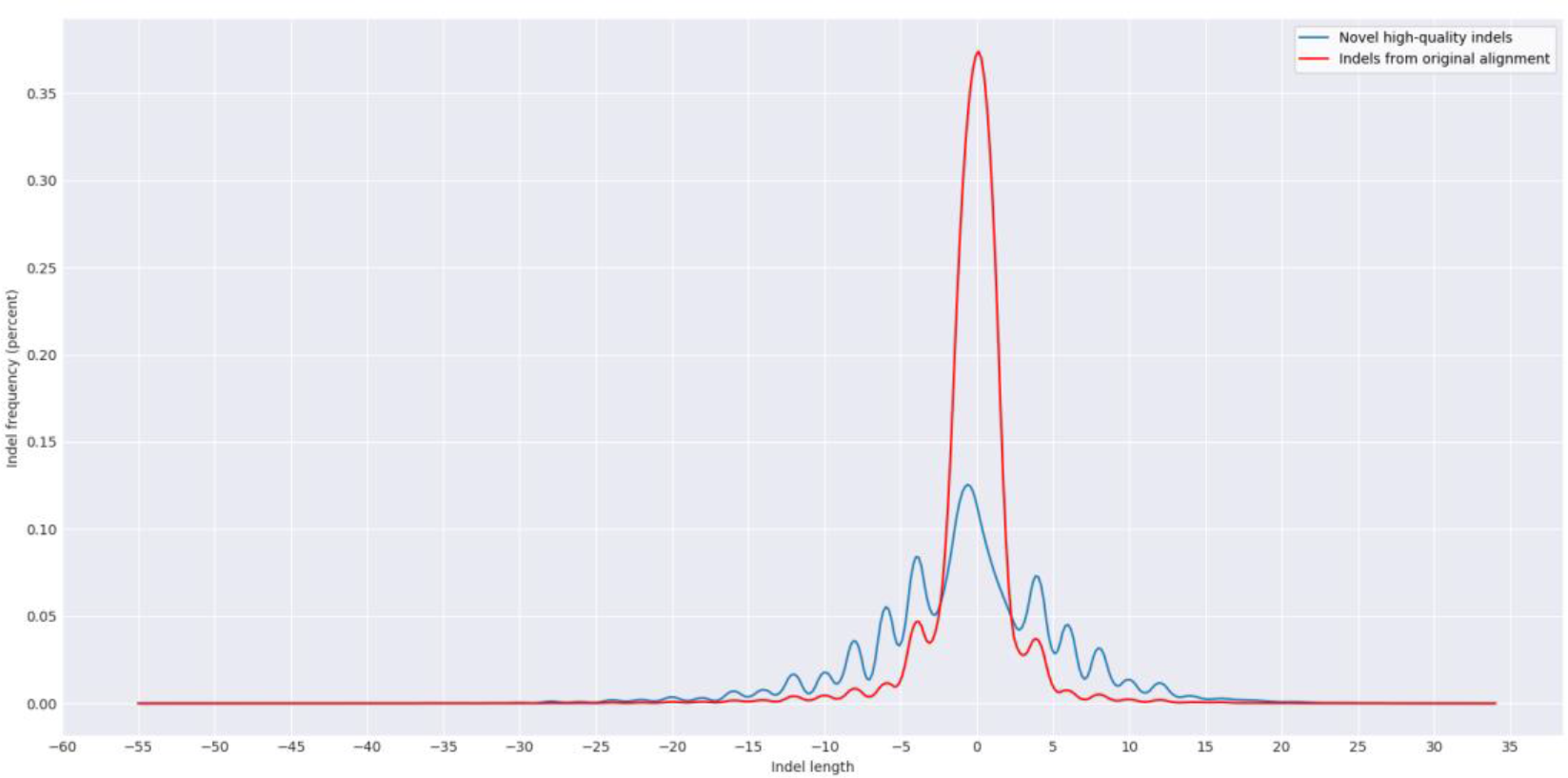
Length Distribution of the novel high-quality indels and indels from the originally mapped reads for all samples. Here a negative value indicates the deletion length.

### Newly mapped reads can add more support to indels not recognized in the originally mapped reads

Most variant calling programs rely on hard evidence for indels marked in the alignment and therefore require a minimum number of reads to support an indel. This step is required to distinguish real variants from the artifacts of sequencing errors. As shown in Figure 8, a 9-base deletion cannot be called from the original alignment due to lack of read support (Figure 8, upper panel). After the mapping of the quality controlled originally unmapped reads, such indels get enough read support and hence are called by the variant caller (Figure 8, lower panel). This explains one of the reasons why these indels are initially missed but got rescued by leveraging the unmapped reads.

**Figure 8:**
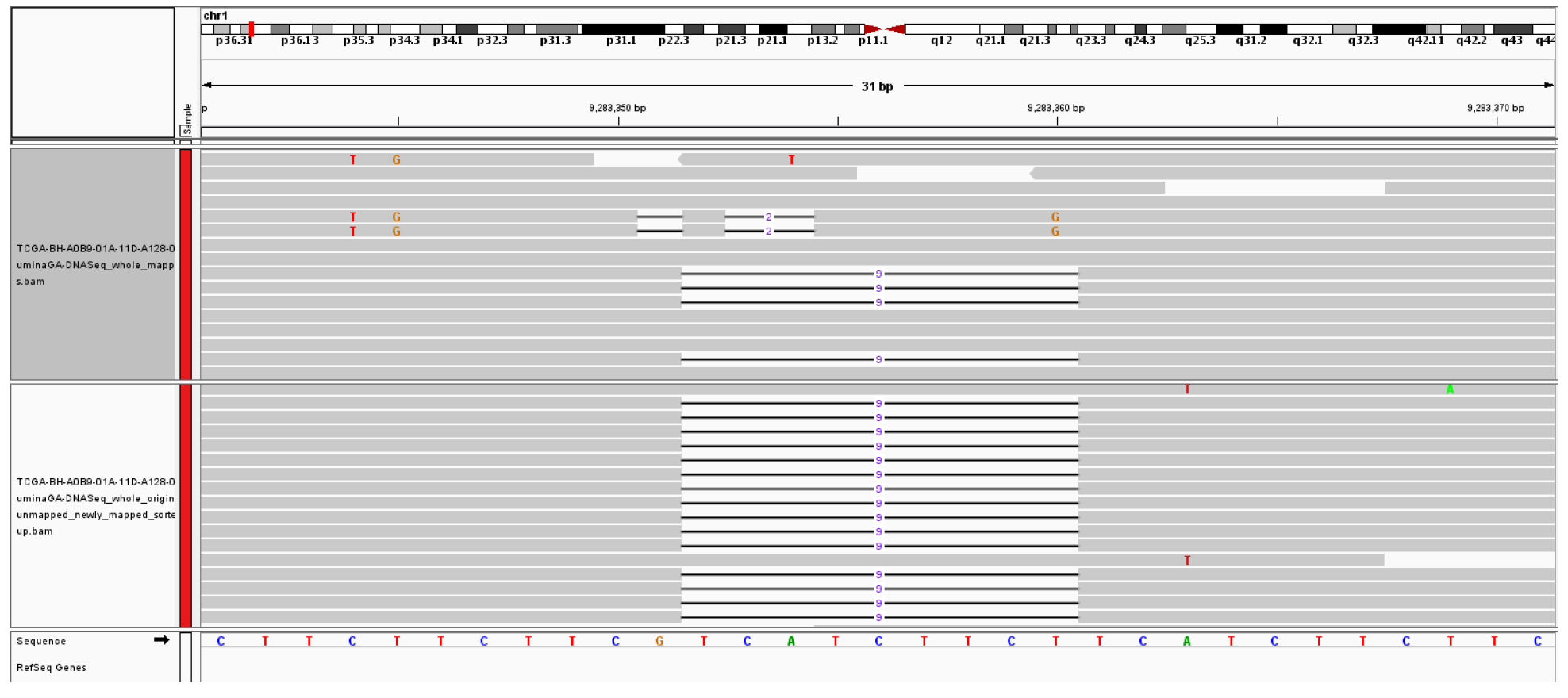
An example of a 9-base deletion which is not initially called from the original alignment due to lack of read-support but later called after the mapping of originally unmapped reads. Here the upper panel shows the original alignment and lower panel shows the alignment of the newly mapped reads.

### Clustering of the samples based on quality-controlled unmapped reads

The samples are compared pairwise using the quality controlled unmapped reads in order to identify biologically relevant signals and to cluster the samples based on the number of similar reads. Pairwise distance is calculated for the unmapped reads from each sample using Mash, a distance estimator based on MinHash [38]. This pairwise distance is then used to cluster the samples. Figure 9 shows the hierarchical clustering of the samples based on Mash distance. The clustering results are then compared with the samples’ PAM50 subtypes collected from TCGA [39]. Out of the total 30 samples, 16 belong to the Basal subtype and the remaining 14 belong to LumA. As shown in figure 9, all but three samples (samples 12, 13, and 14) cluster with the samples of their respective subtype. These three samples belong to Basal subtype but clustered with the samples from LumA subtype. The result reveals that the unmapped reads are most commonly shared among the samples of the same subtype and suggests that these unmapped reads might contribute to the divergence between the two subtypes investigated here. This result also implies that perhaps there is a subtype-specific common cause of mapping failure. These results indicate that the sets of unmapped reads contain sequence information specific to sample subtype and hence leveraging such information may help understand or interpret the related biological questions.

**Figure 9:**
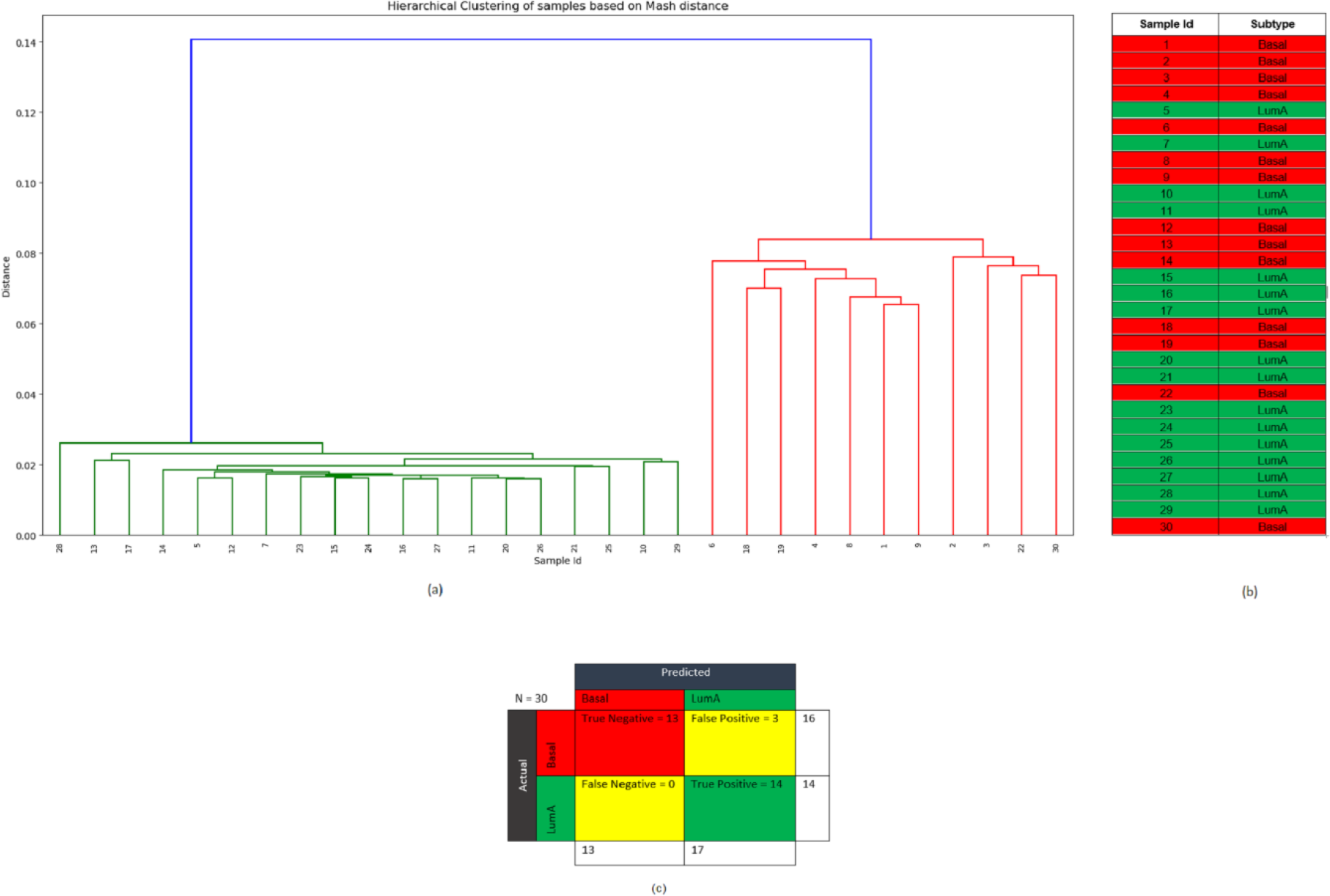
(a) Hierarchical clustering of the samples based on the pairwise Mash distance of the unmapped reads from each sample. (b) PAM50 subtype of the samples from TCGA. Here the Red color corresponds to Basal and green color corresponds to LumA subtype. (c) The confusion matrix.

### Subtype-specific indels from the novel high-quality indels

Indels in each of the samples are counted and all but three samples (samples 12, 13, and 14) in the Basal samples contain 5,818 indels on average and the samples that belong to LumA subtype contain 473 indels, on average. Samples 12, 13, and 14 contain 484, 415, and 535 indels, respectively, i.e., close to the number of indels found in the LumA samples. Similar phenomena are also observed for these three samples in the indels from the originally mapped reads, giving more evidence that these samples actually belong to LumA subtype but mislabeled as Basal, consistent with the result of clustering based on unmapped reads. In addition, the number of newly mapped reads in Samples 12, 13, and 14 are 3.6, 2.5, and 3.5 million, respectively which is closer to the number of newly mapped reads in LumA samples (average number of newly mapped reads = 3.3 million) than to the basal samples (average number of newly mapped reads = 37.89 million). This suggests possible subtype mislabeling of these samples.

Novel high-quality indels are checked to see if they are specific to any of the two subtypes (Basal and LumA) investigated here. An indel is defined as specific to a subtype when it is found in the samples of one subtype and not in the samples of the other subtype. After checking all of the novel high-quality indels, it is found that 89 of the indels are specific to Basal subtype and no such indels are found from the samples investigated here for the LumA subtype.

### Novel high-quality indels overlapped with the oncogenes and tumor suppressor genes

To see which oncogenes and tumor suppressor genes are frequently affected by the newly discovered indels, a list consisting of 142 protein-coding genes (79 oncogenes and 63 tumor suppressor genes) (see Methods) is overlapped with the novel high-quality indels using BEDtools [40]. In total, 62 out of these 142 genes overlapped with these indels. Among these 62 genes, 32 are oncogenes and the remaining ones are tumor suppressor genes. Table 1 lists the top ten genes with the number of indels identified in these genes. The highest number of indels (54) are identified in RUNX1 (Runt Related Transcription Factor 1), a protein-coding tumor suppressor gene. A gene with that many unique indels definitely contains the important signature of breast cancer. Although RUNX1 has received attention as a gene fusion in acute myeloid leukemia (AML) [41, 42], a putative link to breast cancer has recently emerged [43]. However, this gene has not gained enough attention and hence its role in breast cancer remains elusive [44]. One reason for the understudy of the RUNX1 gene is the underpowered expression profile studies as identified by Janes et al. [45]. Another reason, as the result shows here, could be because of not discovering the indels hidden in the unmapped reads. Therefore this study is complementary to the study performed by Janes et al. [45].

**Table 1:** Top ten oncogene and tumor suppressor genes and the number of indels identified in these genes.

| Gene | Number of Indel |
| --- | --- |
| RUNX1 | 54 |
| SYK | 13 |
| CBLB | 11 |
| ETV4 | 11 |
| CCND3 | 10 |
| ETV6 | 9 |
| MAML2 | 7 |
| PIK3CA | 7 |
| BMPR1A | 6 |
| EGFR | 5 |

Frameshift indels are more abundant than in-frame indels in both oncogenes and tumor suppressor genes. Some genes contain only in-frame indel and some contain only frameshift indels. As shown in Table 2, out of the 62 genes overlapped with the novel high-quality indels, 46 contain either in-frame or frameshift indels. The remaining 16 genes contain both in-frame and frameshift indels. As shown in the previous section, frameshift indels are the dominant type of indels and RUNX1 contains the maximum number of indels for both in-frame (12) and frameshift (42) among all genes investigated here, making it an important gene in breast cancer progression.

**Table 2:** List of oncogenes and tumor suppressor genes containing either in-frame or frameshift novel high-quality indels.

| Indel Type | Gene |
| --- | --- |
| In-frame | Oncogenes (5): HMGA2, TPR, RAF1, ROS1, FGFR2 |
|  | Tumor suppressor genes (3): EXT1, CREB1, GPC3 |
| Frameshift | Oncogenes (19): ATF1, CBLB, AKT2, LMO2, TET2, ETV4, BCR, MAF, SMO, PPARG, CARD11, DDX6, PLAG1, EGFR, ABL1, NTRK1, BCL11A, BCL2, FGFR1 |
|  | Tumor suppressor genes (19): RB1, SMARCB1, FLT3, BRCA1, CDH1, SUFU, CHEK2, ARHGEF12, FBXW7, MSH2, NUP98, SUZ12, NPM1, BCL11B, IDH1, EXT2, NR4A3, ATM, BMPR1A |

### Novel high-quality indels mostly alter cancer-related genes than noncancer-related genes

A list of whole genome protein-coding genes not containing the oncogene and tumor suppressor gene is generated from the whole genome gene list produced by GENCODE (version 28 lift37). This list contains 20,172 genes and among these, 6,829 genes are found overlapped with the novel high-quality indels. A hypothesis testing is done to compare the proportion of novel high-quality indels appearing in cancer-related genes (oncogene and tumor suppressor gene) to that in noncancer genes. Statistically, the hypothesis being tested is as follows,

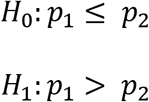

where *p*_1_ denotes the proportion of oncogenes and tumor suppressor genes that overlapped with the novel high-quality indels, i.e., 62/142 = 0.44 and *p*_2_ denotes the proportion of noncancer genes that overlapped with the novel high-quality indels, i.e., 6829/20172 = 0.34.

To test this hypothesis, the standard normal test statistic, *z* is calculated as follows.

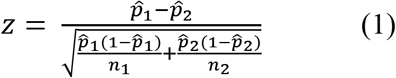

Considering the tail area, α = 0.05, the critical point for this right-tailed test is *z*_0.05_ = 1.64. Based on these data and using the formula (1), the observed value of z is 2.35. Since the value of z exceeds the critical point of 1.64, the null hypothesis, *H*_0_: *p*_1_ ≤ *p*_2_ can be rejected at the α = 0.05 (p-value = 0.009386706). Therefore, the proportion of cancer genes overlapping with novel high-quality indels is significantly higher than the proportion for noncancer genes.

A permutation test can also be done to see if the novel high-quality indels have a higher enrichment in cancer genes (oncogenes and tumor suppressor genes) than in noncancer genes. For this test, from the list containing 20,172 genes (the whole genome gene list not containing the oncogene and tumor suppressor genes), 142 genes (number of total oncogene and tumor suppressor genes) are sampled randomly and repeatedly. The bootstrapping is done 1,000 times yielding 1,000 sets containing 142 genes each.

**Figure 10:**
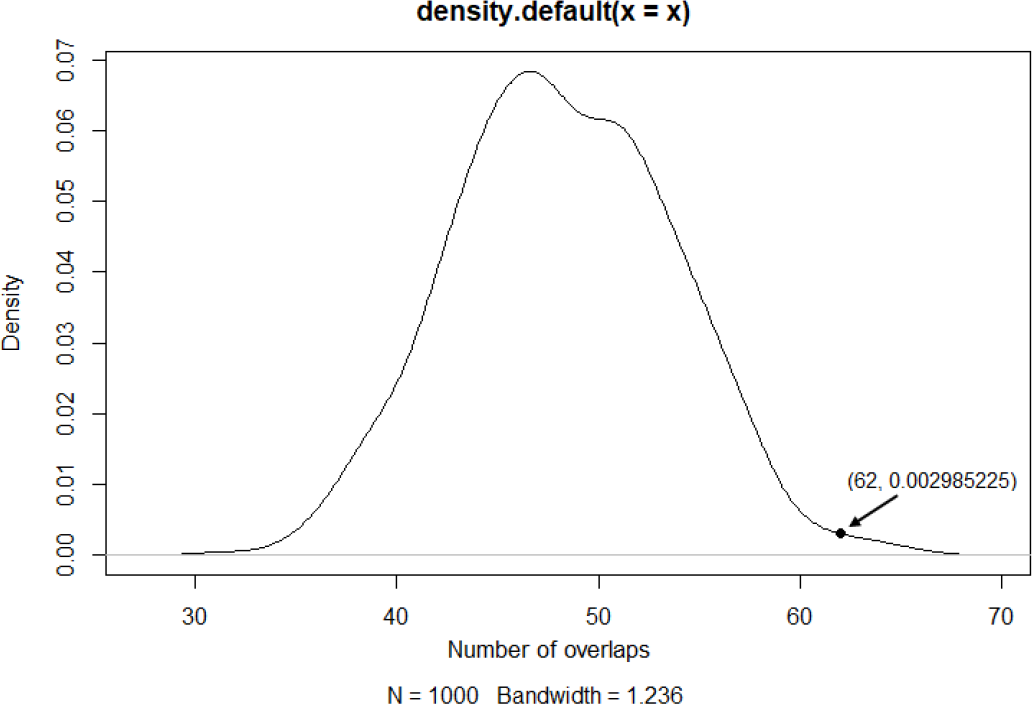
Probability density plot for 1,000 sets containing randomly selected 142 genes overlapped with the novel high-quality indels. The black dot on the right tail represents the value for 62, the number of cancer genes (oncogenes and tumor suppressor genes) overlapped with the novel high-quality indels.

To test the hypothesis, each of these 1,000 sets of randomly picked 142 genes is overlapped with the novel high-quality indels to yield a null distribution of the number of overlaps. Figure 10 shows the probability density plot of such overlap count for 1,000 repetitions. As determined in the previous section, the number of cancer genes (oncogenes and tumor suppressor genes) overlapped with novel high-quality indel is 62. Therefore, the probability density for the number of overlap > 62 (right tail of the density plot) in these random set of indels is calculated as the p-value of the one-sided permutation test. Only in 9 out of 1,000 sets, the number of overlap is higher than 62 and therefore, the p-value is 9/1000 = 0.009 (which is also consistent with the p-value from the z-test above) and since it is < 0.05, the null hypothesis can be rejected. Therefore, it can be concluded that the enrichment of the novel high-quality indels in cancer genes is significantly higher than that in noncancer genes.

### Annotating the novel high-quality indels using Variant Effect Predictor (VEP)

For this analysis, the novel high-quality indels are annotated using Variant Effect Predictor (VEP) [46]. 16,141 of the indels are identified as novel, i.e., not annotated in the Ensembl variation database consisting of dbSNP, Cancer Gene Census, ClinVar, COSMIC, dbGap, DGVa etc. This indicates the significance of this study in rescuing these novel high-quality indels from the discarded reads that can potentially be annotated.

The combined list of indels overlapped with 15,229 genes, 32,335 transcripts, and 2,136 regulatory features. As shown in Figure 11(a), around 75% of the novel high-quality indels are in the noncoding regions located in the intron or intergenic region. Figure 11(b) shows that 72% of the indels are frameshift indels which cause a disruption of the translational reading frame and can have a disruptive impact in the protein by causing protein truncation and/or loss of function. In addition, a small amount of the indels are “Splice donor variants” changing the 2-base region at the 5′ end of an intron and can have a similar impact as frameshift indels. 25% of the indels are in-frame indels having a “moderate” impact in the protein by not disrupting the protein but changing the effectiveness of that protein. 70% of the novel-high quality indels having disruptive and moderate impact overlap with the protein-coding transcripts, the leading biotype of all features (Transcripts, Regulatory Features, and Motif Features) as shown in Figure 11(c). Out of the remaining indels, 37.44% of the indels (12.48% of the total novel high-quality indels) have a “modifier” impact overlap with long intergenic RNA transcripts (lincRNA). LincRNAs are noncoding transcripts with a length longer than 200 nucleotides and are the largest class of noncoding RNA molecules in the human genome. There is emerging evidence that noncoding RNAs regulate gene expression by influencing chromatin modification, mRNA splicing, and protein translation [47, 48] as well as contribute to mammary tumor development [49, 50] and progression. Therefore the novel high-quality indels overlapped with these transcripts deserve more attention and studying these indels has biological significance.

**Figure 11:**
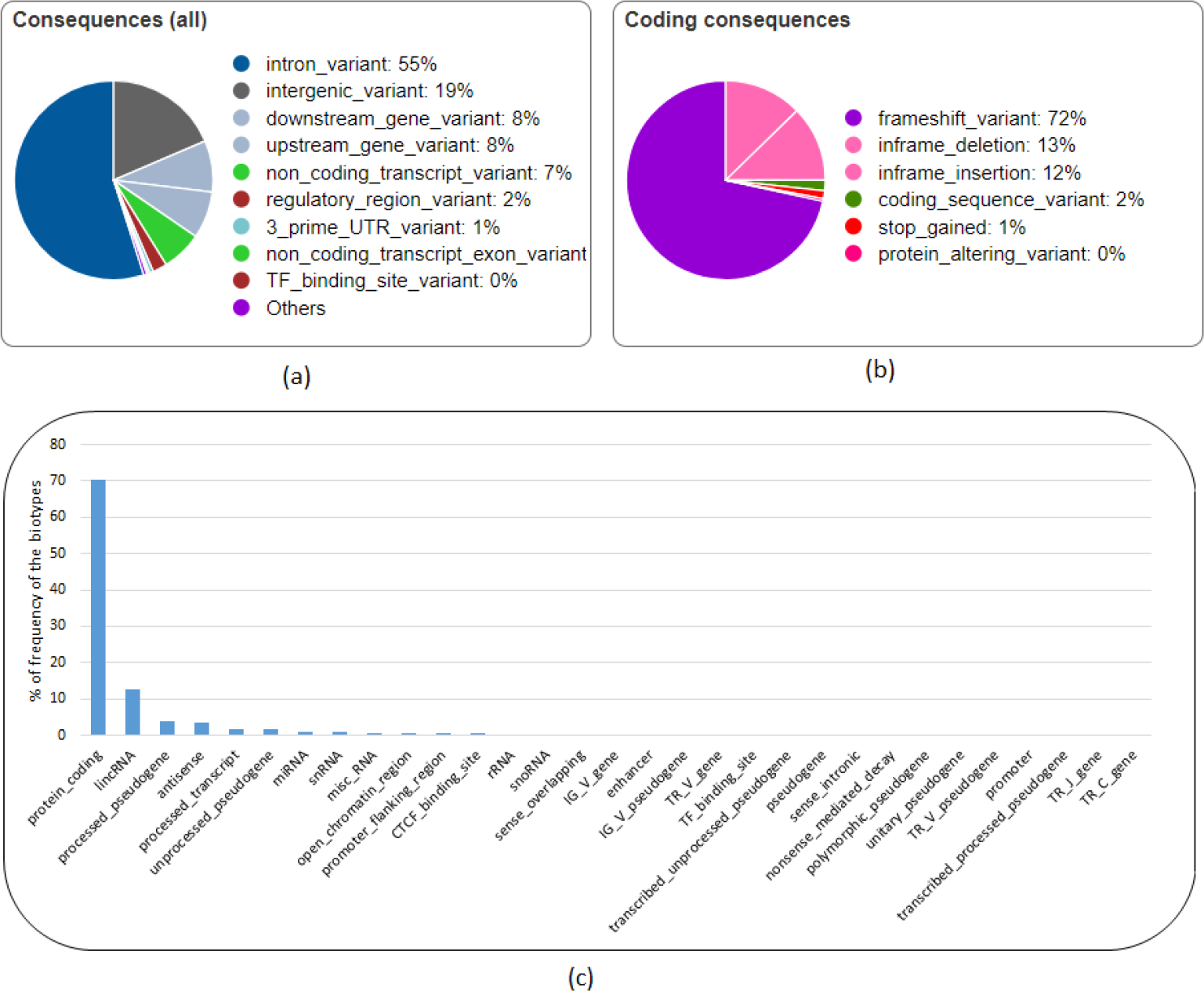
Analysis of the novel high-quality indels using Variant Effect Predictor (VEP). (a) All consequences (b) coding consequences, (c) distribution of biotype of the features overlapped with the novel high-quality indels.

### Functional annotation of the overlapping genes

Functional annotation of the genes overlapping with the novel high-quality indels using David [51] (version 6.7) shows strong correlation with “Pathways in Cancer” (Fisher Exact p-value: 6.2 × 10^−5^), “PI3K-Akt signaling pathway” (Fisher Exact p-value: 1.5 × 10^−4^), RAP1 signaling pathway (Fisher Exact p-value: 10^−4^), and RAS signaling pathway (Fisher Exact p-value: 4.7 × 10^−3^). A previous study shows that components of the PI3K-Akt signaling pathway are recurrently altered in cancers and the survival signals induced by several receptors are mediated mainly by this pathway [52]. Ras-associated protein-1 (RAP1) is an important regulator of cell functions and has been found playing a vital role in cell invasion and metastasis in cancers [53]. The signaling pathways involving RAS protein can contribute to tumor growth, survival, and spread, and play a crucial role in the pathogenesis of other hematologic malignancies as well [54, 55]. Therefore, these results suggest that the newly found indels interacting with these genes may participate in cancer-related biological processes and play an important role in cancer progression.

### Genes missed in the original mapping but found in novel high-quality indels show association with cancer and other diseases

There are 42 genes overlapping with the novel high-quality indels but not with the indels from the originally mapped reads. Table 3 lists the genes with their types. Functional annotation of the protein-coding genes using David shows that these genes are related to biological process such as immune response, protein localization, protein transport, regulation of transcription, and regulation of RNA metabolic process which can control molecular functions such as antigen binding, peptide binding, MHC protein binding, and peptide-antigen binding. In addition, these genes are associated with protein domain such as Immunoglobulin subtype and Krueppel-Associated Box (KRAB)-Zinc Finger Protein (ZFP). Immunoglobulin subtype is involved in cell-cell recognition, cell-surface receptors, muscle structure, and the immune system [56] and therapy targeting this protein domain has been used for liver cancer [57], breast cancer [58], and Follicular Lymphoma [59, 60]. Krueppel-Associated Box (KRAB)-Zinc Finger Protein (ZFP) is the largest class of transcription factors in the human genome [61] and is largely involved in tumorigenesis [62].

**Table 3:** Name and type of the genes that overlap with the novel high-quality indels but with the indels from the originally mapped reads.

| Gene Type | Gene Name |
| --- | --- |
| **Antisense** | RP11-534L20.5, JMJD1C-AS1, CTB-22K21.2, RP11-1079K10.4, RP11-16N11.2, RP11-26P13.2, RP11-1299A16.3, RP1-16A9.1. |
| **LincRNA** | RP5-1065P14.2, RP11-309G3.3, RP11-382D12.1, RP11-14C22.3, RP11-386I8.6, LINC00379, RP3-503A6.2, RP3-416J7.4, RP11-100L22.4. |
| **miRNA** | MIR4477A. |
| **Pseudogene** | RP5-857K21.7, RP11-428G5.7, BNIP3P1, RP11-713H12.2, VN1R90P, MLLT10P1, UNC93B3, TBCAP3, CTD-2158P22.1, RP11-823P9.3, RP3- |
|  | 416J7.1, CYP4F44P, OR5BH1P, PABPC1P3, CASKP1, TRBV12-1, TRBV12-2. |
| **Protein coding** | OR10G8, ZNF26, ZNF84, KIF20A. |
| **snRNA** | RNU1-59P, RNU6-377P, RNU1-36P. |

A PubMed search returned results for three genes namely KIF20A, BNIP3P1, and ZNF84.

Kinesin family member 20A (KIF20A), also known as RAB6KIFL, is a member of the kinesin superfamily of motor proteins, a conserved motor domain which binds to microtubules to generate the energy required for trafficking of proteins and organelles during the growth of numerous cancers [63, 64]. KIF20A is found overexpressed at both the mRNA and protein levels than the normal counterparts in breast cancer [65–67] and also in several other cancers including gastric cancer [68], bladder cancer [69, 70], pancreatic cancer [71–73], hepatocellular cancer [74], lung cancer [75], glioma [76], and melanoma [77]. The overexpression of KIF20A gene is significantly associated with the poor survival of breast cancer patient [65, 66] and drug resistance [66, 78]. Similar phenomena are observed with other cancer patients as well [68, 70, 71, 73, 75, 79]. Silencing or knockdown of KIF20A can significantly inhibit cell proliferation and cancer progression [72, 80]. Therefore, KIF20A has been suggested as a direct therapeutic target [72, 81] and KIF20A-derived peptide has been used in immunotherapy in clinical trials to improve the prognosis of cancer patients [63, 76, 82–85]. Although KIF20A has a strong association with breast cancer, no mutation is found in this gene from the originally mapped reads which shows the limitation of the current approach and this limitation can be alleviated by exploring the unmapped reads. Besides cancer, KIF20A is found associated with heart disease in infants. A recent study by Louw et al. [86] identified an undescribed type of lethal congenital restrictive cardiomyopathy, a disease affecting the right ventricle of two siblings. Exome sequencing analysis of these affected siblings and their unaffected sibling revealed two compound heterozygous variants in KIF20A; a maternal missense variant (c.544C>T: p. R182W) changing an arginine to a tryptophan and a paternal frameshift deletion (c.1905delT: p. S635Tfs15, in exon 15) that introduces a premature stop codon 15 amino acids downstream. Louw et al.[86] validated the variants by Sanger sequencing, found the presence of both variants in the affected siblings, and confirmed a heterozygous carrier status in both parents. In addition, both variants were absent in the unaffected sibling. The C>T missense SNP does not let KIF20A support efficient transport of Aurora B as part of the chromosomal passenger complex causing Aurora B trapped on chromatin during the cell division and hence it fails to translocate to the spindle midzone during cytokinesis. This claim is verified by Louw et al. [86] in the zebrafish model where translational blocking of KIF20A resulted in a cardiomyopathy phenotype. A similar congenital restrictive cardiomyopathy is also identified to be caused by the deletion resulting in loss-of-function of KIF20A [86]. Despite such significance, these two variants that affect protein function were absent in the population control exome such as ExAC Browser database, a catalog of genetic data of 60,706 humans of various ethnicities [87]. The missense variant was found in two individuals from South Asia and Europe and the frameshift deletion was present in 32 individuals of African descent [86, 88]. This observation supports the claim that clinically important mutations can be missed and one of the reasons might be because of overlooking the unmapped reads. By exploring the unmapped read of 30 breast cancer patients, Genesis-indel finds a frameshift deletion of T (chr5:137520225 CT -> C) that overlaps with the exon of KIF20A gene (Figure 12).

**Figure 12:**
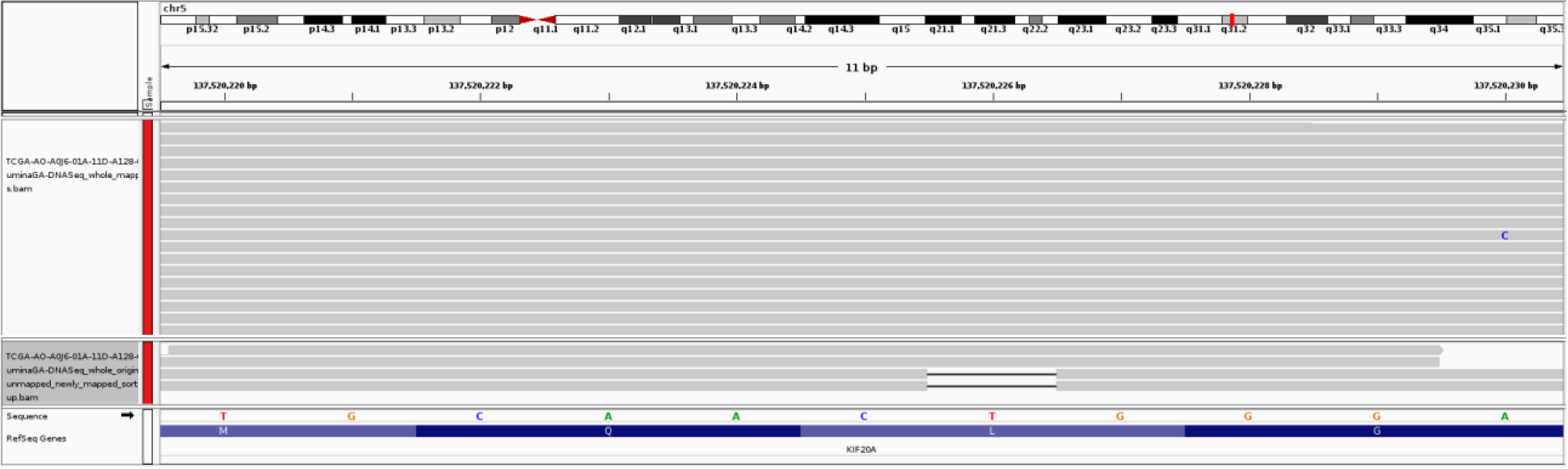
**A** 1-base deletion in the exon of KIF20A gene which is not initially called from the original alignment but called after the mapping of originally unmapped reads. The upper panel shows the original alignment and lower panel shows the alignment of the newly mapped reads.

Among the remaining genes, BCL2 interacting protein 3 pseudogene 1 (BNIP3P1) is found to be upregulated in patients with Breast Cancer Brain Metastases when compared to Breast Cancer (76% vs. 24%) or compared to Primary Brain Tumors (74% vs. 26%) [89] and is suggested to be used as a molecular biomarker for Breast Cancer Brain Metastases. Zinc Finger Protein 84 (ZNF84) is found significantly associated with tumor size and TNM (Tumor, Node, Metastases) staging for cervical cancer and squamous cell carcinoma and in vitro validation shows that it promotes cell proliferation via AKT signaling pathway [90]. Although the literature doesn’t show any association between these genes and breast cancer, it is worth exploring due to their association with other cancers.

Out of the 42 genes, two LincRNAs namely RP3-416J7.4 and RP11-386I8.6 contain the same number of indels as the protein-coding genes. Although little is known about their association with breast cancer, analysis using TANRIC [91] on TCGA-BRCA data reveals that these two LincRNAs are differentially expressed (t-test p-value = 0.000023337 and 0.003812, respectively) between the carriers and non-carriers of somatic mutations in the TP53 gene, a tumor suppressor gene spontaneously found altered in breast carcinomas [92].

While this paper shows the significance of uncovering novel high-quality indels from the originally unmapped reads in patients with breast cancer, there are a couple of limitations. Firstly, this study is conducted by using a computational pipeline. Though the pipeline is computationally feasible and results are convincing as well as supported by experimentally validated literature, it lacks some validation experiments in vivo to govern the clinical importance of the newly identified indels. Secondly, filtering indels solely based on the “PASS” flag may cause missing rare variants. Therefore, an algorithm such as ForestQC [93] which combines traditional variant filtering approach with machine learning algorithm to determine the quality of the variant can be incorporated to the present pipeline to improve the quality control procedure and achieve better results.

## Conclusion

This paper emphasizes the interest of the unmapped reads and the potential loss of important information and describes Genesis-indel, a computational pipeline to rescue novel high-quality indels by exploring unmapped reads that are normally discarded from the downstream analysis.

Analyzing the whole genome DNA alignment of 30 breast cancer patients from TCGA reveals that there are a nonnegligible number of unmapped reads that are overlooked earlier. These unmapped reads can be a reservoir of important biological information and can affect the variant identification. Pairwise comparisons of the unmapped reads from the samples highlights that the unmapped reads are most commonly shared among the samples of the same subtype and might contribute to the divergence between subtypes. In addition, it indicates that the unmapped reads contain sequence information specific to sample subtypes and there might be a subtype-specific common cause of mapping failure. After the quality control, 65.38% of the originally unmapped reads from 30 breast cancer patients got mapped to the reference genome. Genesis-indel finds 72,997 novel high-quality indels of diverse lengths and 16,141 of these indels have not been annotated in any of the variation database used by Ensembl. This finding will enable these indels to be subjected to further study. These novel high-quality indels are mainly enriched in frameshift indels and have high to moderate impact in the protein. These indels mostly alter the oncogenes and tumor suppressor genes and overlap with genes significantly related to different cancer pathways. Moreover, these indels overlap with genes not found in the indels from the originally mapped reads and functional annotation shows that these genes contribute to the development and growth of tumor in multiple carcinomas. Therefore, these findings collectively suggest that complete characterization of these indels is essential for downstream cancer research. Genesis-indel is expected to be highly useful for uncovering the missed indels that can be further explored for clinical decision making.

## Methods

### The Genesis-indel pipeline

Genesis-indel is designed to leverage unmapped reads from an alignment with the goal to rescue indels that are hidden in the discarded unmapped reads. Figure 2 shows the schematic of the Genesis-indel workflow. The input to Genesis-indel is the alignment file (BAM file) of the patient genome and the reference genome. In the preprocessing step, Genesis-indel extracts the unmapped alignment by checking the alignment flag using SAMtools (version 1.4) [24]. From this, it extracts the “Originally Unmapped” reads using SAMtools and stores the reads in a FASTQ file. This FASTQ file is then processed by Trimmomatic (version 0.36) [26] to do the quality control of the unmapped reads by removing adapter sequences. In this experiment, the Illumina adapter, TruSeq2 for single-end reads are removed. Moreover, low quality or N bases where the base quality is below 3 are removed from both ends of the reads (LEADING:3, TRAILING:3). Reads are scanned with a 4-base wide sliding window and reads are cut when the average quality per base drops below 15 (SLIDINGWINDOW: 4:15). Reads with length below the 36 bases are dropped (MINLEN:36). These quality controlled single-end reads are used as input to the mapper in the next step.

The quality controlled unmapped reads are mapped to the reference genome using BWA-MEM (version 0.7.15-r1140) [28], a sensitive mapper to map reads with indels. After the reads are aligned by BWA-MEM, some reads still remain unmapped. These reads are aligned to the reference genome using BLAT (BLAST-Like Alignment Tool) [29], another sensitive local alignment tool. At the end of this step, the alignments from BWA-MEM and BLAT are merged. The resultant alignment is sorted and indexed using SAMtools and duplicates are marked in the newly mapped reads using MarkDuplicates tool from Picard (version 1.65) [94]. After read alignment and duplicate removal, indels are called using Platypus (version 0.7.9.1). Separately, indels are also called from the original (input) BAM file. Indels found only in the newly mapped reads and not in the original alignment are reported as novel indels. After identifying the novel indels, another step of filtering is done to keep only the high-quality indels, i.e., the indels that are called with high confidence by Platypus. Therefore, only the indels with the “PASS” flags are reported at the final step. These are the novel high-quality indels reported in the final output and selected for downstream analysis.

### Preparing a list of oncogene and tumor suppressor genes

A list of oncogenes and tumor suppressor genes is obtained from an online resource [95], a list compiled from the CancerGenes [96]. While preparing the list, if a gene is marked as both an oncogene and a tumor suppressor gene in CancerGenes, a literature search is performed to determine the gene’s role in tumor development. Any gene with an ambiguous role as an oncogene or tumor suppressor gene is excluded from the list. The final list contains 79 oncogenes and 63 tumor suppressor genes. The start and end positions of the genes are obtained from GENCODE (version 28 lift37). Supplementary Table S1 contains the list of the genes and their positions.

### Software availability and system requirements

Genesis-indel is implemented in C++ and can run on any operating systems that have a C++ compiler. The source code and the command line version of Genesis-indel are freely available at https://github.com/mshabbirhasan/Genesis-indel. Users are welcome to report bugs and provide comments through the issue tracker on GitHub. The README and GitHub wiki describes the command line options available in Genesis-indel with examples. Although Genesis-indel uses BWA-MEM as the mapper and Platypus as the default variant caller, future version will allow the user the flexibility to customize the program and use the mapper and caller of their choice by doing little modification in the script. Default Genesis-indel usually takes 24 hours to process a sample with 50× coverage in a workstation with an intel core i7 CPU and 128 GB of RAM.

### Data availability

No new data sample is generated for this study. The alignment file (BAM) for the 30 breast cancer patients are obtained from The Cancer Genome Atlas (TCGA) project ((https://portal.gdc.cancer.gov/). Supplementary Table S2 lists the TCGA Sample Barcode and alignment filename for the patients. The reference genome used is Homo_sapiens_assembly19.fasta, the same reference used by TCGA to align the reads. The annotation of the genes is collected from GENCODE (version 28 lift37). All other data supporting the findings of this study are available within this article and in the supplementary materials. These data are also available from the authors upon request.

## Supporting information

## Acknowledgments

This work is partially supported by Virginia Tech’s Open Access Subvention Fund. Authors acknowledge The Cancer Genome Atlas (TCGA) (http://cancergenome.nih.gov) as the primary source of data. Authors thank Gustavo Arango and Saima Tithi from ZhangLab at Virginia Tech for helpful discussions and feedback.

## Author contributions

M.S.H. and L.Z. conceptualized and designed the research. M.S.H. developed and tested the Genesis-indel pipeline and performed data analysis. M.S.H and X.W. did the statistical analysis. L.Z. supervised the research. All authors wrote and approved the manuscript.

## Reference

1. MacArthur DG, Tyler-Smith C: Loss-of-function variants in the genomes of healthy humans. Human molecular genetics 2010, 19(R2):R125–R130.

2. Paschka P, Marcucci G, Ruppert AS, Mrózek K, Chen H, Kittles RA, Vukosavljevic T, Perrotti D, Vardiman JW, Carroll AJ: Adverse prognostic significance of KIT mutations in adult acute myeloid leukemia with inv (16) and t (8; 21): a Cancer and Leukemia Group B Study. Journal of Clinical Oncology 2006, 24(24):3904–3911.

3. Sequist LV, Martins RG, Spigel D, Grunberg SM, Spira A, Jänne PA, Joshi VA, McCollum D, Evans TL, Muzikansky A: First-line gefitinib in patients with advanced non-small-cell lung cancer harboring somatic EGFR mutations. Journal of clinical oncology 2008, 26(15):2442–2449.

4. Li H, Ruan J, Durbin R: Mapping short DNA sequencing reads and calling variants using mapping quality scores. Genome research 2008, 18(11):1851–1858.

5. Li R, Li Y, Kristiansen K, Wang J: SOAP: short oligonucleotide alignment program. Bioinformatics 2008, 24(5):713–714.

6. Li H, Durbin R: Fast and accurate short read alignment with Burrows-Wheeler transform. Bioinformatics 2009, 25(14):1754–1760.

7. Langmead B, Trapnell C, Pop M, Salzberg SL: Ultrafast and memory-efficient alignment of short DNA sequences to the human genome. Genome biology 2009, 10(3):R25.

8. Langmead B, Salzberg SL: Fast gapped-read alignment with Bowtie 2. Nature methods 2012, 9(4):357.

9. Zaharia M, Bolosky WJ, Curtis K, Fox A, Patterson D, Shenker S, Stoica I, Karp RM, Sittler T: Faster and more accurate sequence alignment with SNAP. arXiv preprint arXiv:11115572 2011.

10. Li R, Yu C, Li Y, Lam T-W, Yiu S-M, Kristiansen K, Wang J: SOAP2: an improved ultrafast tool for short read alignment. Bioinformatics 2009, 25(15):1966–1967.

11. Koboldt DC, Ding L, Mardis ER, Wilson RK: Challenges of sequencing human genomes. Briefings in bioinformatics 2010, 11(5):484–498.

12. Mitsudomi T, Yatabe Y: Epidermal growth factor receptor in relation to tumor development: EGFR gene and cancer. The FEBS journal 2010, 277(2):301–308.

13. Yasuda H, Kobayashi S, Costa DB: EGFR exon 20 insertion mutations in non-small-cell lung cancer: preclinical data and clinical implications. The lancet oncology 2012, 13(1):e23–e31.

14. Bhangale TR, Rieder MJ, Livingston RJ, Nickerson DA: Comprehensive identification and characterization of diallelic insertion-deletion polymorphisms in 330 human candidate genes. Human molecular genetics 2005, 14(1):59–69.

15. Dawson E, Chen Y, Hunt S, Smink LJ, Hunt A, Rice K, Livingston S, Bumpstead S, Bruskiewich R, Sham P: A SNP resource for human chromosome 22: extracting dense clusters of SNPs from the genomic sequence. Genome research 2001, 11(1):170–178.

16. Mullaney JM, Mills RE, Pittard WS, Devine SE: Small insertions and deletions (INDELs) in human genomes. Human molecular genetics 2010, 19(R2):R131–R136.

17. Collins FS, Drumm ML, Cole JL, Lockwood WK, Woude GV, Iannuzzi MC: Construction of a general human chromosome jumping library, with application to cystic fibrosis. Science 1987, 235(4792):1046–1049.

18. Warren ST, Zhang F, Licameli GR, Peters JF: The fragile X site in somatic cell hybrids: an approach for molecular cloning of fragile sites. Science 1987, 237(4813):420–423.

19. Falini B, Mecucci C, Tiacci E, Alcalay M, Rosati R, Pasqualucci L, La Starza R, Diverio D, Colombo E, Santucci A: Cytoplasmic nucleophosmin in acute myelogenous leukemia with a normal karyotype. New England Journal of Medicine 2005, 352(3):254–266.

20. Nakao M, Yokota S, Iwai T, Kaneko H, Horiike S, Kashima K, Sonoda Y, Fujimoto T, Misawa S: Internal tandem duplication of the flt3 gene found in acute myeloid leukemia. Leukemia 1996, 10(12):1911–1918.

21. Sequist LV, Martins RG, Spigel D, Grunberg SM, Spira A, Jänne PA, Joshi VA, McCollum D, Evans TL, Muzikansky A: First-line gefitinib in patients with advanced non-small-cell lung cancer harboring somatic EGFR mutations. Journal of Clinical Oncology 2008, 26(15):2442–2449.

22. Cheung VG, Spielman RS: Genetics of human gene expression: mapping DNA variants that influence gene expression. Nature Reviews Genetics 2009, 10(9):595–604.

23. Weinstein JN, Collisson EA, Mills GB, Shaw KRM, Ozenberger BA, Ellrott K, Shmulevich I, Sander C, Stuart JM, Network CGAR: The cancer genome atlas pancancer analysis project. Nature genetics 2013, 45(10):1113.

24. Li H, Handsaker B, Wysoker A, Fennell T, Ruan J, Homer N, Marth G, Abecasis G, Durbin R: The sequence alignment/map format and SAMtools. Bioinformatics 2009, 25(16):2078–2079.

25. Andrews S: FastQC: a quality control tool for high throughput sequence data. 2010.

26. Bolger AM, Lohse M, Usadel B: Trimmomatic: a flexible trimmer for Illumina sequence data. Bioinformatics 2014, 30(15):2114–2120.

27. Peng X, Wang J, Zhang Z, Xiao Q, Li M, Pan Y: Re-alignment of the unmapped reads with base quality score. In: Bmc Bioinformatics: 2015. BioMed Central: S8.

28. Li H: Aligning sequence reads, clone sequences and assembly contigs with BWA-MEM. arXivpreprint arXiv:13033997 2013.

29. Kent WJ: BLAT—the BLAST-like alignment tool. Genome research 2002, 12(4):656–664.

30. Rimmer A, Phan H, Mathieson I, Iqbal Z, Twigg SR, Wilkie AO, McVean G, Lunter G, Consortium W: Integrating mapping-, assembly-and haplotype-based approaches for calling variants in clinical sequencing applications. Nature genetics 2014, 46(8):912.

31. Hasan MS, Wu X, Zhang L: Performance evaluation of indel calling tools using real short-read data. Human genomics 2015, 9(1):20.

32. Thorvaldsdóttir H, Robinson JT, Mesirov JP: Integrative Genomics Viewer (IGV): high-performance genomics data visualization and exploration. Briefings in bioinformatics 2013, 14(2):178–192.

33. Rampino N, Yamamoto H, Ionov Y, Li Y, Sawai H, Reed JC, Perucho M: Somatic frameshift mutations in the BAX gene in colon cancers of the microsatellite mutator phenotype. Science 1997, 275(5302):967–969.

34. Li J, Yen C, Liaw D, Podsypanina K, Bose S, Wang SI, Puc J, Miliaresis C, Rodgers L, McCombie R: PTEN, a putative protein tyrosine phosphatase gene mutated in human brain, breast, and prostate cancer. science 1997, 275(5308):1943–1947.

35. Ogura Y, Bonen DK, Inohara N, Nicolae DL, Chen FF, Ramos R, Britton H, Moran T, Karaliuskas R, Duerr RH: A frameshift mutation in NOD2 associated with susceptibility to Crohn’s disease. Nature 2001, 411(6837):603.

36. Iengar P: An analysis of substitution, deletion and insertion mutations in cancer genes. Nucleic acids research 2012, 40(14):6401–6413.

37. Mills RE, Pittard WS, Mullaney JM, Farooq U, Creasy TH, Mahurkar AA, Kemeza DM, Strassler DS, Ponting CP, Webber C: Natural genetic variation caused by small insertions and deletions in the human genome. Genome research 2011:gr. 115907.115110.

38. Ondov BD, Treangen TJ, Melsted P, Mallonee AB, Bergman NH, Koren S, Phillippy AM: Mash: fast genome and metagenome distance estimation using MinHash. Genome biology 2016, 17(1):132.

39. Network CGA: Comprehensive molecular portraits of human breast tumours. Nature 2012, 490(7418):61.

40. Quinlan AR, Hall IM: BEDTools: a flexible suite of utilities for comparing genomic features. Bioinformatics 2010, 26(6):841–842.

41. Silva FP, Morolli B, Storlazzi CT, Anelli L, Wessels H, Bezrookove V, Kluin-Nelemans HC, Giphart-Gassler M: Identification of RUNX1/AML1 as a classical tumor suppressor gene. Oncogene 2003, 22(4):538.

42. Miyoshi H, Shimizu K, Kozu T, Maseki N, Kaneko Y, Ohki M: t (8; 21) breakpoints on chromosome 21 in acute myeloid leukemia are clustered within a limited region of a single gene, AML1. Proceedings of the National Academy of Sciences 1991, 88(23):10431–10434.

43. Ferrari N, Mohammed ZM, Nixon C, Mason SM, Mallon E, McMillan DC, Morris JS, Cameron ER, Edwards J, Blyth K: Expression of RUNX1 correlates with poor patient prognosis in triple negative breast cancer. PloS one 2014, 9(6):e100759.

44. Browne G, Taipaleenmäki H, Bishop NM, Madasu SC, Shaw LM, Van Wijnen AJ, Stein JL, Stein GS, Lian JB: Runx1 is associated with breast cancer progression in MMTV-PyMT transgenic mice and its depletion in vitro inhibits migration and invasion. Journal of cellular physiology 2015, 230(10):2522–2532.

45. Janes KA: RUNX1 and its understudied role in breast cancer. Cell cycle 2011, 10(20):3461–3465.

46. McLaren W, Gil L, Hunt SE, Riat HS, Ritchie GR, Thormann A, Flicek P, Cunningham F: The ensembl variant effect predictor. Genome biology 2016, 17(1):122.

47. Rinn JL, Chang HY: Genome regulation by long noncoding RNAs. Annual review of biochemistry 2012, 81:145–166.

48. Roberts TC, Morris KV, Weinberg MS: Perspectives on the mechanism of transcriptional regulation by long non-coding RNAs. Epigenetics 2014, 9(1):13–20.

49. Silva JM, Boczek NJ, Berres MW, Ma X, Smith DI: LSINCT5 is over expressed in breast and ovarian cancer and affects cellular proliferation. RNA biology 2011, 8(3):496–505.

50. Gupta RA, Shah N, Wang KC, Kim J, Horlings HM, Wong DJ, Tsai M-C, Hung T, Argani P, Rinn JL: Long non-coding RNA HOTAIR reprograms chromatin state to promote cancer metastasis. Nature 2010, 464(7291):1071.

51. Dennis G, Sherman BT, Hosack DA, Yang J, Gao W, Lane HC, Lempicki RA: DAVID: database for annotation, visualization, and integrated discovery. Genome biology 2003, 4(9):R60.

52. Fresno JV, Casado E, Cejas P, Belda-Iniesta C, González-Barón M: PI3K/Akt signalling pathway and cancer. Cancer treatment reviews 2004, 30(2):193–204.

53. Zhang Y-L, Wang R-C, Cheng K, Ring BZ, Su L: Roles of Rap1 signaling in tumor cell migration and invasion. Cancer biology & medicine 2017, 14(1):90.

54. Downward J: Targeting RAS signalling pathways in cancer therapy. Nature Reviews Cancer 2003, 3(1):11.

55. Reuter CW, Morgan MA, Bergmann L: Targeting the Ras signaling pathway: a rational, mechanism-based treatment for hematologic malignancies? Blood 2000, 96(5):1655–1669.

56. Teichmann SA, Chothia C: Immunoglobulin superfamily proteins in Caenorhabditis elegans. Journal of molecular biology 2000, 296(5):1367–1383.

57. Ettinger D, Order S, Wharam M, Parker MK, Klein J, Leichner P: Phase I-II study of isotopic immunoglobulin therapy for primary liver cancer. Cancer treatment reports 1982, 66(2):289–297.

58. Musolino A, Naldi N, Bortesi B, Pezzuolo D, Capelletti M, Missale G, Laccabue D, Zerbini A, Camisa R, Bisagni G: Immunoglobulin G fragment C receptor polymorphisms and clinical efficacy of trastuzumab-based therapy in patients with HER-2/neu-positive metastatic breast cancer. Journal of Clinical Oncology 2008, 26(11):1789–1796.

59. Weng W-K, Levy R: Two immunoglobulin G fragment C receptor polymorphisms independently predict response to rituximab in patients with follicular lymphoma. Journal of clinical oncology 2003, 21(21):3940–3947.

60. Weng W-K, Czerwinski D, Timmerman J, Hsu FJ, Levy R: Clinical outcome of lymphoma patients after idiotype vaccination is correlated with humoral immune response and immunoglobulin G Fc receptor genotype. Journal of clinical oncology 2004, 22(23):4717–4724.

61. Mark C, Abrink M, Hellman L: Comparative analysis of KRAB zinc finger proteins in rodents and man: evidence for several evolutionarily distinct subfamilies of KRAB zinc finger genes. DNA and cell biology 1999, 18(5):381–396.

62. Cheng Y, Geng H, Cheng SH, Liang P, Bai Y, Li J, Srivastava G, Ng MH, Fukagawa T, Wu X: KRAB zinc finger protein ZNF382 is a proapoptotic tumor suppressor that represses multiple oncogenes and is commonly silenced in multiple carcinomas. Cancer research 2010:0008–5472. CAN-0009-4566.

63. Suzuki N, Hazama S, Ueno T, Matsui H, Shindo Y, Iida M, Yoshimura K, Yoshino S, Takeda K, Oka M: A phase I clinical trial of vaccination with KIF20A-derived peptide in combination with gemcitabine for patients with advanced pancreatic cancer. Journal of immunotherapy (Hagerstown, Md: 1997) 2014, 37(1):36.

64. Vale RD, Reese TS, Sheetz MP: Identification of a novel force-generating protein, kinesin, involved in microtubule-based motility. Cell 1985, 42(1):39–50.

65. Zou JX, Duan Z, Wang J, Sokolov A, Xu J, Chen CZ, Li JJ, Chen H-W: Kinesin family deregulation coordinated by bromodomain protein ANCCA and histone methyltransferase MLL for breast cancer cell growth, survival, and tamoxifen resistance. Molecular Cancer Research 2014.

66. Khongkow P, Gomes A, Gong C, Man E, Tsang JW, Zhao F, Monteiro L, Coombes R, Medema R, Khoo U: Paclitaxel targets FOXM1 to regulate KIF20A in mitotic catastrophe and breast cancer paclitaxel resistance. Oncogene 2016, 35(8):990.

67. Groth-Pedersen L, Aits S, Corcelle-Termeau E, Petersen NH, Nylandsted J, Jäättelä M: Identification of cytoskeleton-associated proteins essential for lysosomal stability and survival of human cancer cells. PloSone 2012, 7(10):e45381.

68. Claerhout S, Lim JY, Choi W, Park Y-Y, Kim K, Kim S-B, Lee J-S, Mills GB, Cho JY: Gene expression signature analysis identifies vorinostat as a candidate therapy for gastric cancer. PloS one 2011, 6(9):e24662.

69. Neef R, Grüneberg U, Barr FA: Assay and Functional Properties of Rabkinesin-6/Rab6-KIFL/MKlp2 in Cytokinesis. Methods in enzymology 2005, 403:618–628.

70. Lu Y, Liu P, Wen W, Grubbs CJ, Townsend RR, Malone JP, Lubet RA, You M: Crossspecies comparison of orthologous gene expression in human bladder cancer and carcinogen-induced rodent models. American journal of translational research 2011, 3(1):8.

71. Taniuchi K, Furihata M, Saibara T: KIF20A-mediated RNA granule transport system promotes the invasiveness of pancreatic cancer cells. Neoplasia 2014, 16(12):1082–1093.

72. Stangel D, Erkan M, Buchholz M, Gress T, Michalski C, Raulefs S, Friess H, Kleeff J: Kif20a inhibition reduces migration and invasion of pancreatic cancer cells. journal of surgical research 2015, 197(1):91–100.

73. Imai K, Hirata S, Irie A, Senju S, Ikuta Y, Yokomine K, Harao M, Inoue M, Tomita Y, Tsunoda T: Identification of HLA-A2-restricted CTL epitopes of a novel tumour-associated antigen, KIF20A, overexpressed in pancreatic cancer. British journal of cancer 2011, 104(2):300.

74. Gasnereau I, Boissan M, Margall-Ducos G, Couchy G, Wendum D, Bourgain-Guglielmetti F, Desdouets C, Lacombe M-L, Zucman-Rossi J, Sobczak-Thépot J: KIF20A mRNA and its product MKlp2 are increased during hepatocyte proliferation and hepatocarcinogenesis. The American journal of pathology 2012, 180(1):131–140.

75. Fang H, Yamaguchi R, Liu X, Daigo Y, Yew PY, Tanikawa C, Matsuda K, Imoto S, Miyano S, Nakamura Y: Quantitative T cell repertoire analysis by deep cDNA sequencing of T cell receptor α and β chains using next-generation sequencing (NGS). Oncoimmunology 2014, 3(12):e968467.

76. Saito K, Ohta S, Kawakami Y, Yoshida K, Toda M: Functional analysis of KIF20A, a potential immunotherapeutic target for glioma. Journal of neuro-oncology 2017, 132(1):63–74.

77. Yamashita J, Fukushima S, Jinnin M, Honda N, Makino K, Sakai K, Masuguchi S, Inoue Y, Ihn H: Kinesin family member 20A is a novel melanoma-associated antigen. Acta dermato-venereologica 2012, 92(6):593–597.

78. Bobustuc GC, Kassam AB, Rovin RA, Jeudy S, Smith JS, Isley B, Singh M, Paranjpe A, Srivenugopal KS, Konduri SD: MGMT inhibition in ER positive breast cancer leads to CDC2, TOP2A, AURKB, CDC20, KIF20A, Cyclin A2, Cyclin B2, Cyclin D1, ERα and Survivin inhibition and enhances response to temozolomide. Oncotarget 2018, 9(51):29727.

79. Ho JR, Chapeaublanc E, Kirkwood L, Nicolle R, Benhamou S, Lebret T, Allory Y, Southgate J, Radvanyi F, Goud B: Deregulation of Rab and Rab effector genes in bladder cancer. PloS one 2012, 7(6):e39469.

80. Taniuchi K, Nakagawa H, Nakamura T, Eguchi H, Ohigashi H, Ishikawa O, Katagiri T, Nakamura Y: Down-regulation of RAB6KIFL/KIF20A, a kinesin involved with membrane trafficking of discs large homologue 5, can attenuate growth of pancreatic cancer cell. Cancer research 2005, 65(1):105–112.

81. Zhang W, He W, Shi Y, Gu H, Li M, Liu Z, Feng Y, Zheng N, Xie C, Zhang Y: High expression of KIF20A is associated with poor overall survival and tumor progression in early-stage cervical squamous cell carcinoma. PloS one 2016, 11(12):e0167449.

82. Asahara S, Takeda K, Yamao K, Maguchi H, Yamaue H: Phase I/II clinical trial using HLA-A24-restricted peptide vaccine derived from KIF20A for patients with advanced pancreatic cancer. Journal of translational medicine 2013, 11(1):291.

83. Aruga A, Takeshita N, Kotera Y, Okuyama R, Matsushita N, Ohta T, Takeda K, Yamamoto M: Phase I clinical trial of multiple-peptide vaccination for patients with advanced biliary tract cancer. Journal of translational medicine 2014, 12(1):61.

84. Fujiwara Y, Okada K, Omori T, Sugimura K, Miyata H, Ohue M, Kobayashi S, Takahashi H, Nakano H, Mochizuki C: Multiple therapeutic peptide vaccines for patients with advanced gastric cancer. International journal of oncology 2017, 50(5):1655–1662.

85. Miyazawa M, Katsuda M, Maguchi H, Katanuma A, Ishii H, Ozaka M, Yamao K, Imaoka H, Kawai M, Hirono S: Phase II clinical trial using novel peptide cocktail vaccine as a postoperative adjuvant treatment for surgically resected pancreatic cancer patients. International journal of cancer 2017, 140(4):973–982.

86. Louw JJ, Bastos RN, Chen X, Verdood C, Corveleyn A, Jia Y, Breckpot J, Gewillig M, Peeters H, Santoro MM: Compound heterozygous loss-of-function mutations in KIF20A are associated with a novel lethal congenital cardiomyopathy in two siblings. PLoS genetics 2018, 14(1):e1007138.

87. Karczewski KJ, Weisburd B, Thomas B, Solomonson M, Ruderfer DM, Kavanagh D, Hamamsy T, Lek M, Samocha KE, Cummings BB: The ExAC browser: displaying reference data information from over 60 000 exomes. Nucleic acids research 2016, 45(D1):D840–D845.

88. Lek M, Karczewski KJ, Minikel EV, Samocha KE, Banks E, Fennell T, O’Donnell-Luria AH, Ware JS, Hill AJ, Cummings BB: Analysis of protein-coding genetic variation in 60,706 humans. Nature 2016, 536(7616):285.

89. Schulten H-J, Bangash M, Karim S, Dallol A, Hussein D, Merdad A, Al-Thoubaity FK, Al-Maghrabi J, Jamal A, Al-Ghamdi F: Comprehensive molecular biomarker identification in breast cancer brain metastases. Journal of translational medicine 2017, 15(1):269.

90. Li P, Guo H, Zhou G, Shi H, Li Z, Guan X, Deng Z, Li S, Zhou S, Wang Y: Increased ZNF84 expression in cervical cancer. Archives of gynecology and obstetrics 2018, 297(6):1525–1532.

91. Li J, Han L, Roebuck P, Diao L, Liu L, Yuan Y, Weinstein JN, Liang H: TANRIC: an interactive open platform to explore the function of lncRNAs in cancer. Cancer research 2015:canres. 0273.2015.

92. Børresen-Dale AL: TP53 and breast cancer. Human mutation 2003, 21(3):292–300.

93. Li J, Jew B, Zhan L, Hwang S, Coppola G, Freimer NB, Sul JH: ForestQC: quality control on genetic variants from next-generation sequencing data using random forest. bioRxiv 2018:444828.

94. Picard [http://broadinstitute.github.io/picard]

95. A list of oncogenes and tumor suppressors used in the comparison of gene functional groups [http://cancerres.aacrjournals.org/content/canres/suppl/2012/01/23/0008-5472.CAN-11-2266.DC1/T374K.pdf]

96. Higgins ME, Claremont M, Major JE, Sander C, Lash AE: CancerGenes: a gene selection resource for cancer genome projects. Nucleic acids research 2006, 35(suppl_1):D721–D726.

