## Supplementary figures and images for "Uncovering missed indels by leveraging unmapped reads"

### supplementary_figure1.pdf

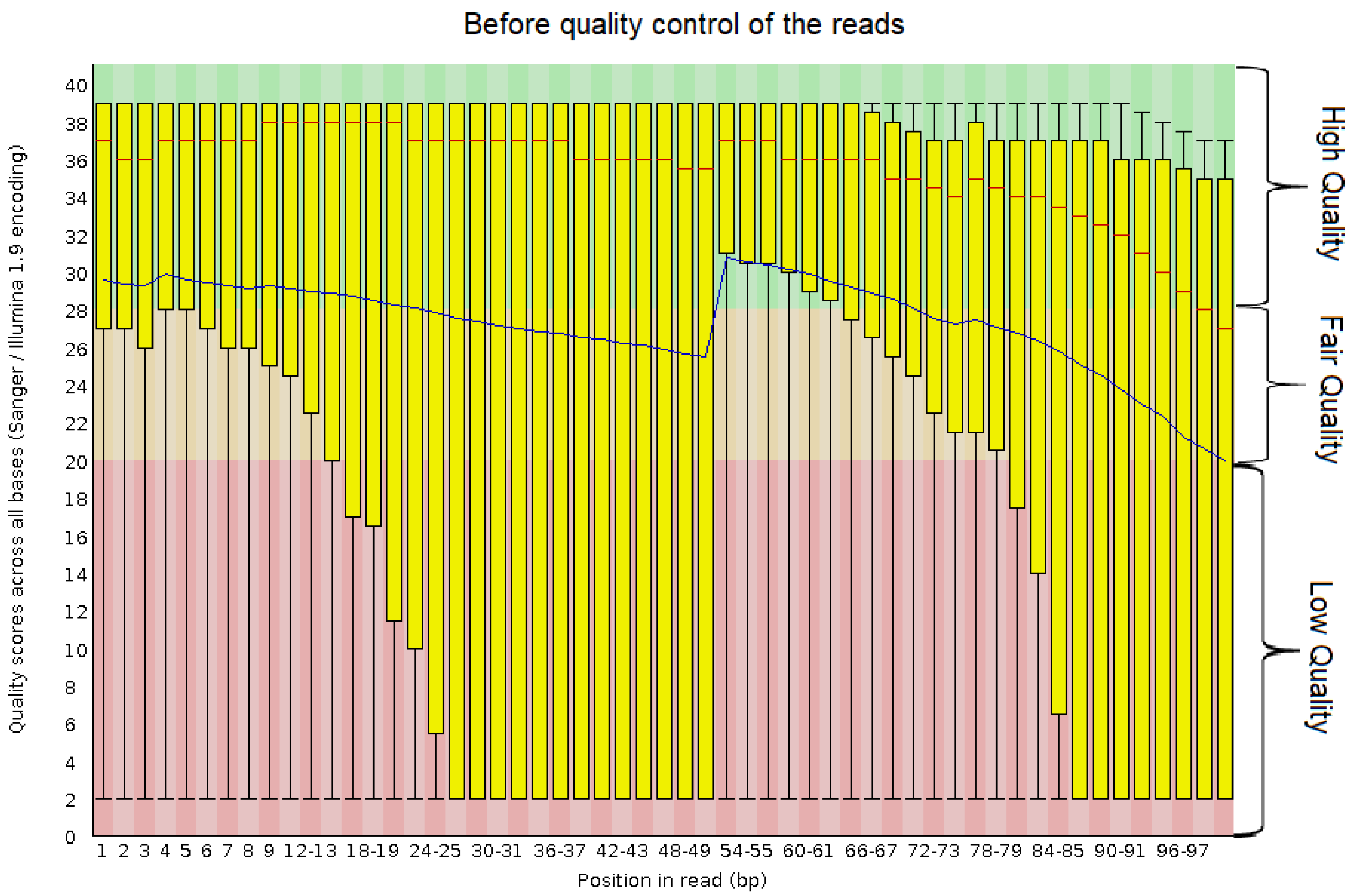

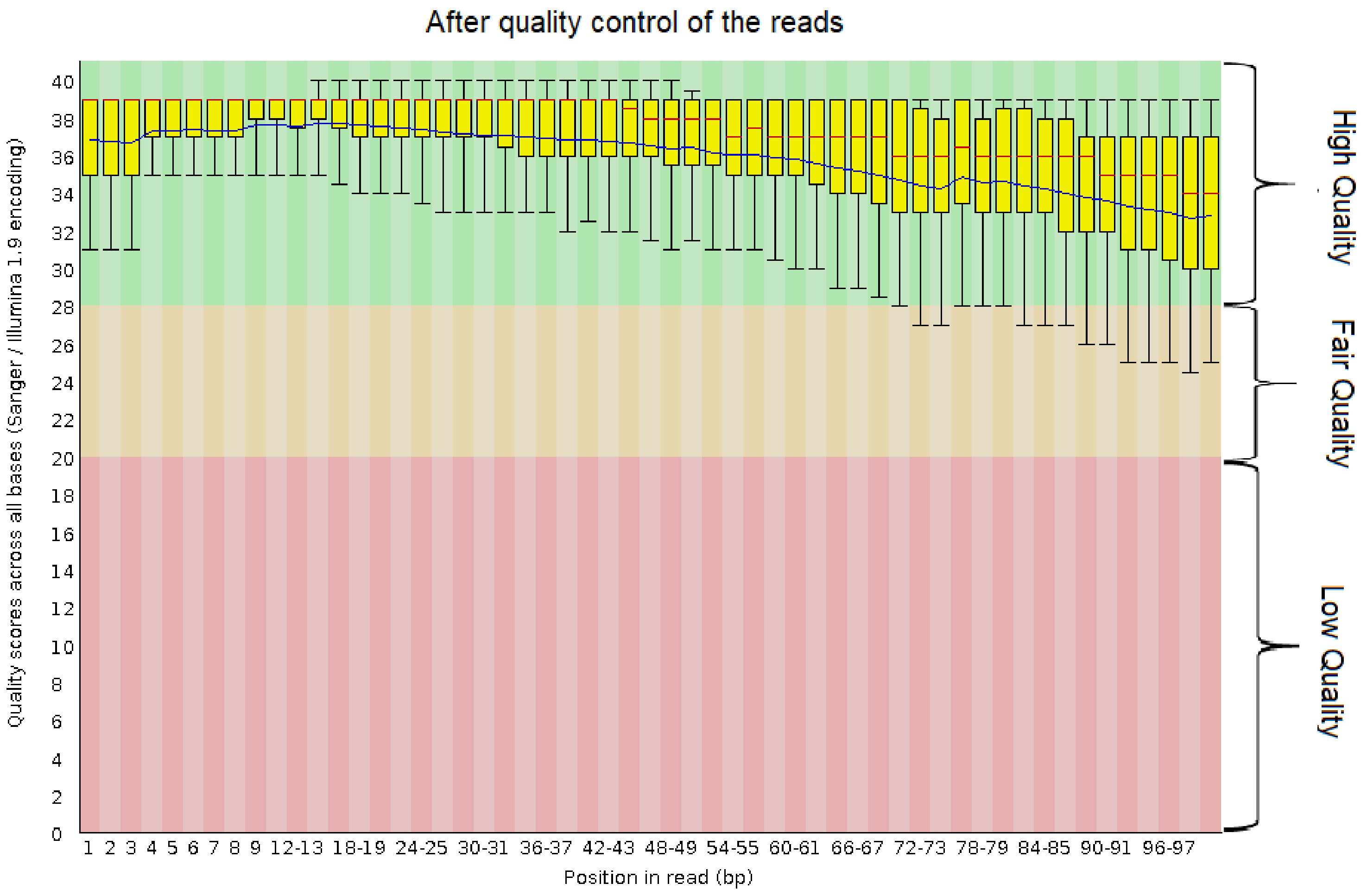

### supplementary_figure2.pdf

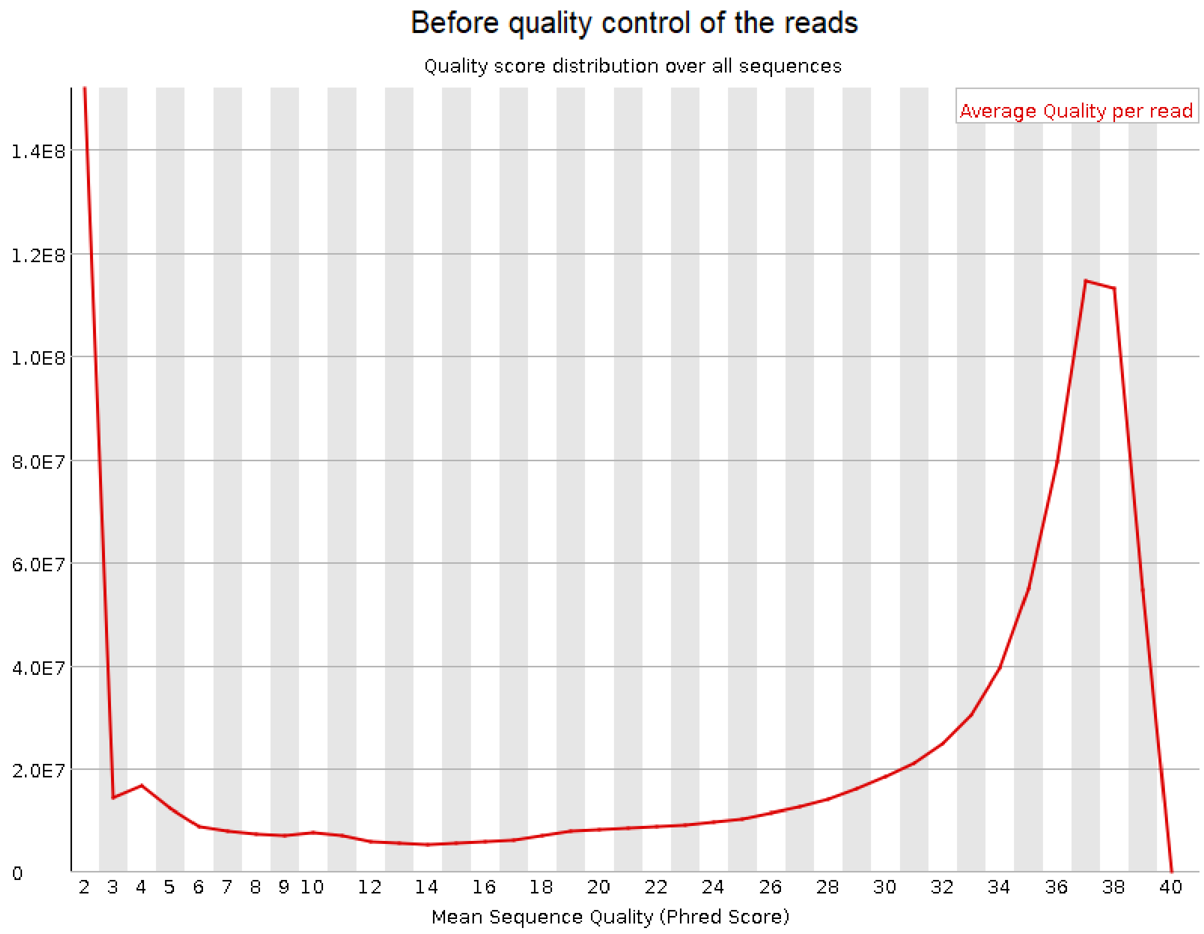

After quality control of the reads

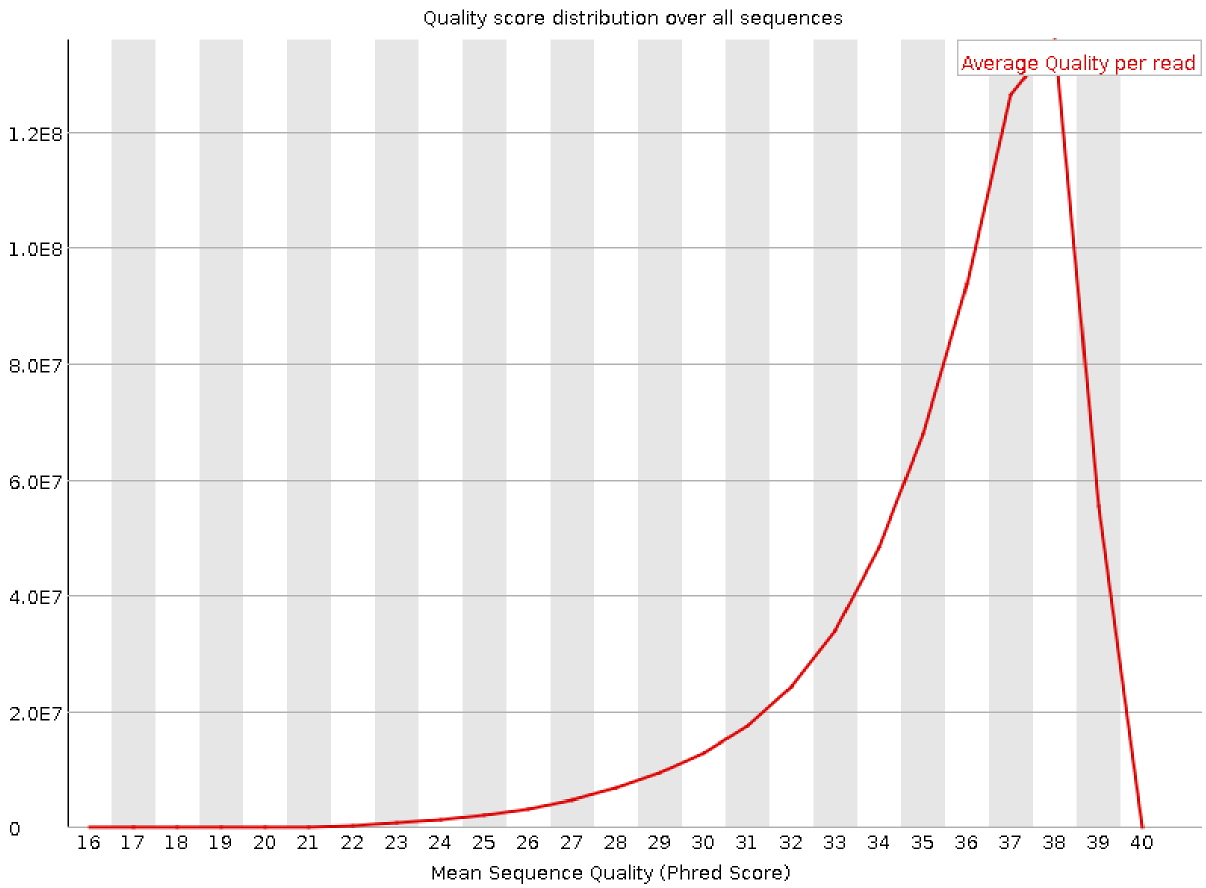


(b)

### supplementary_figure3.pdf

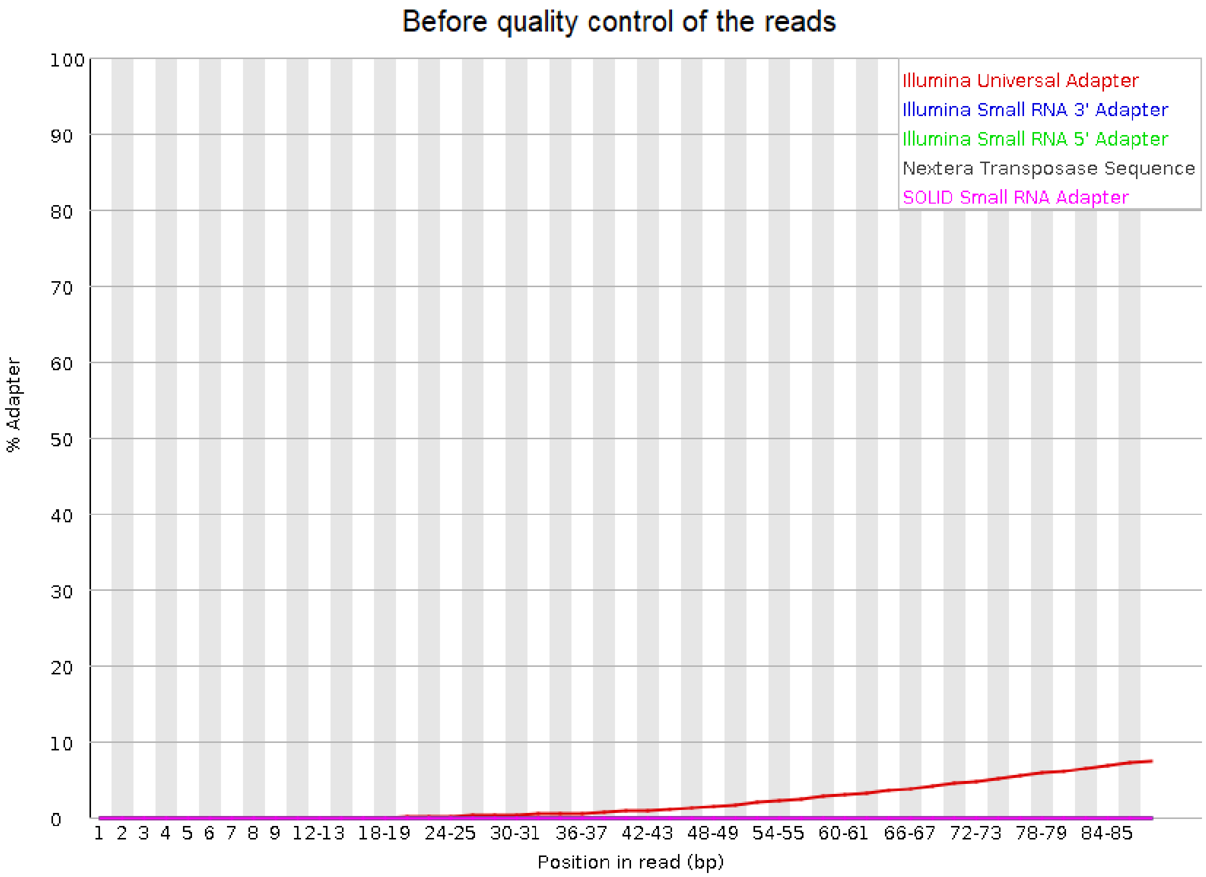

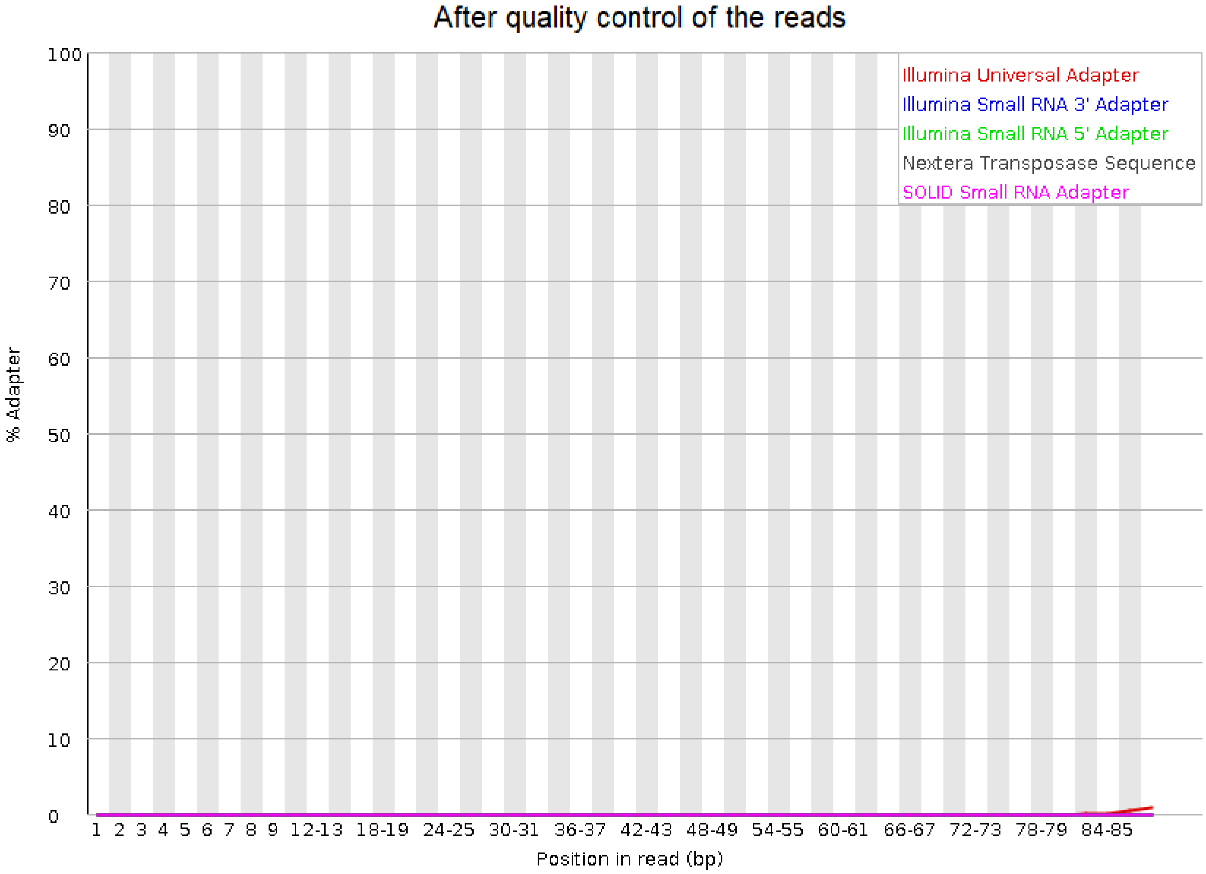

### supplementary_figure4.pdf

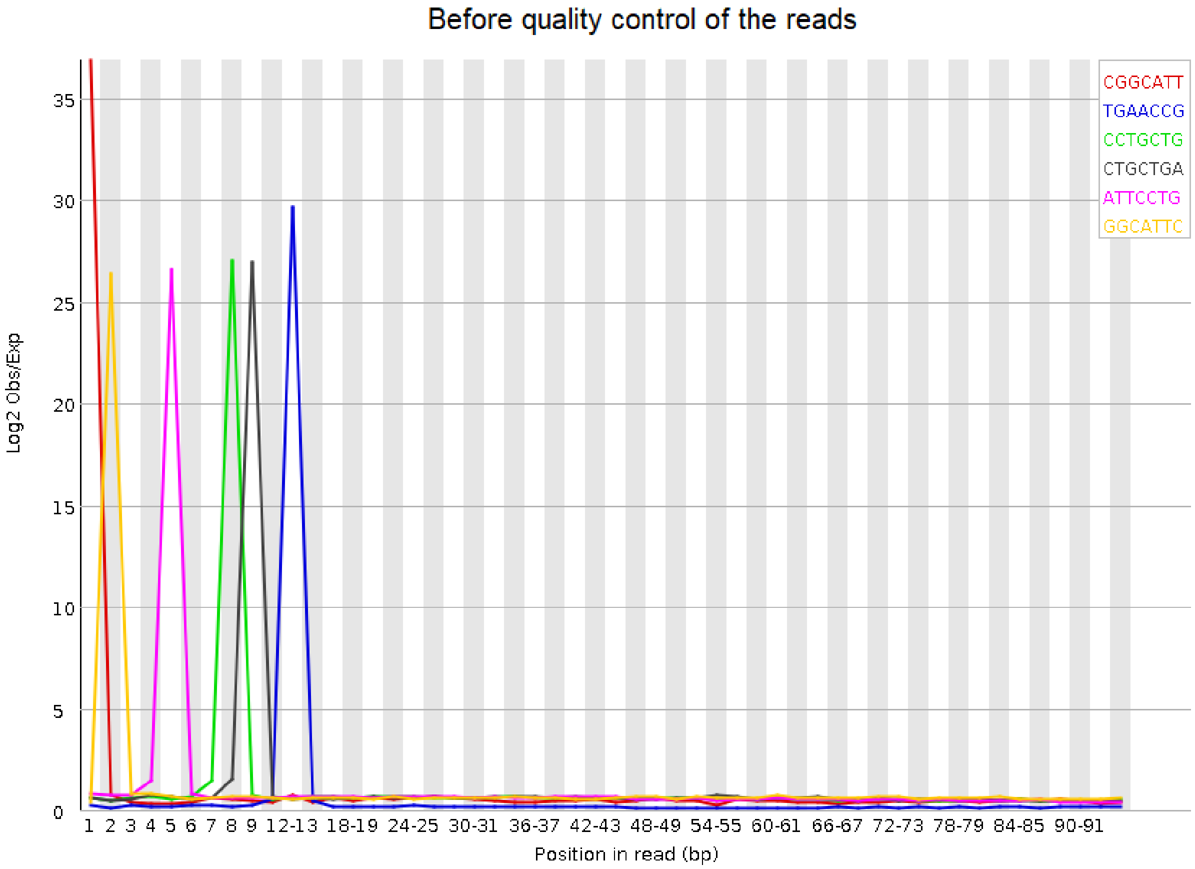

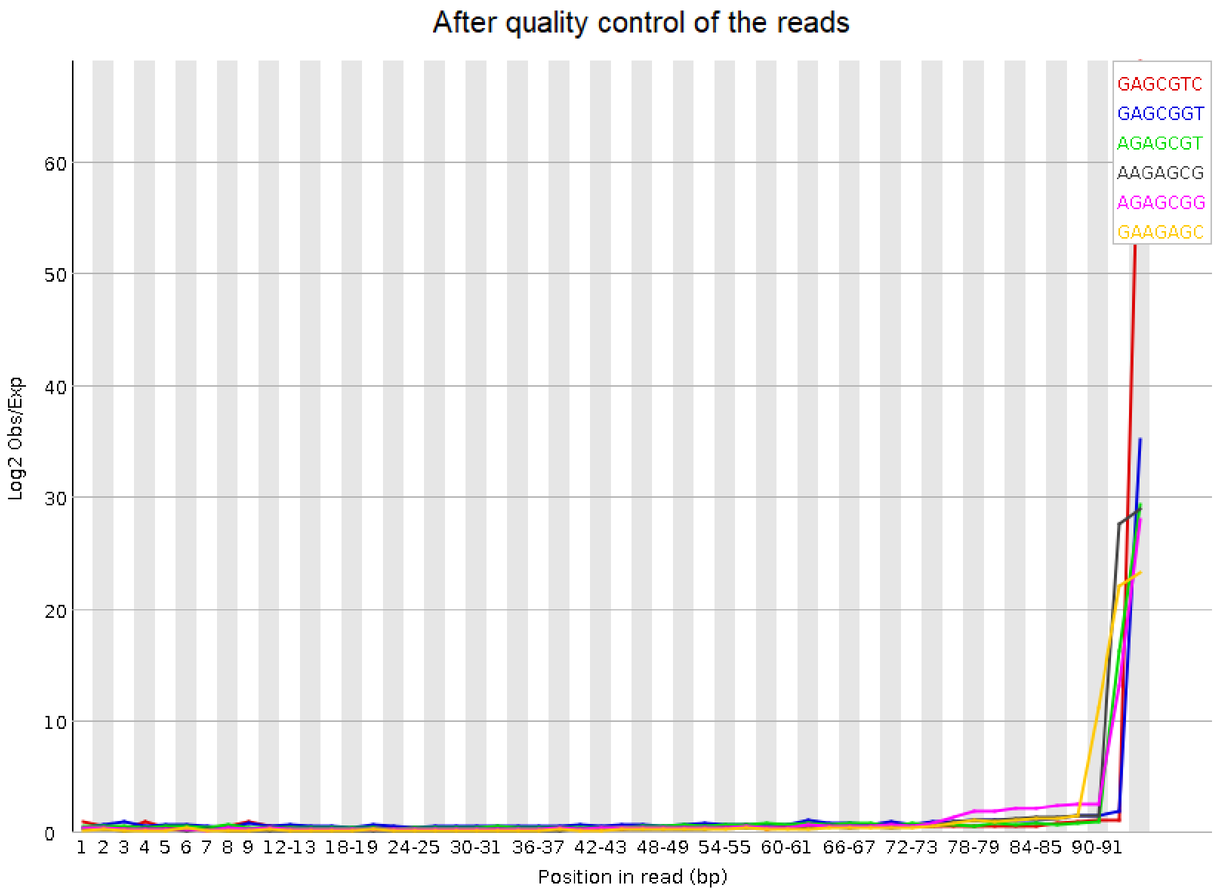

### supplementary_figure5.pdf

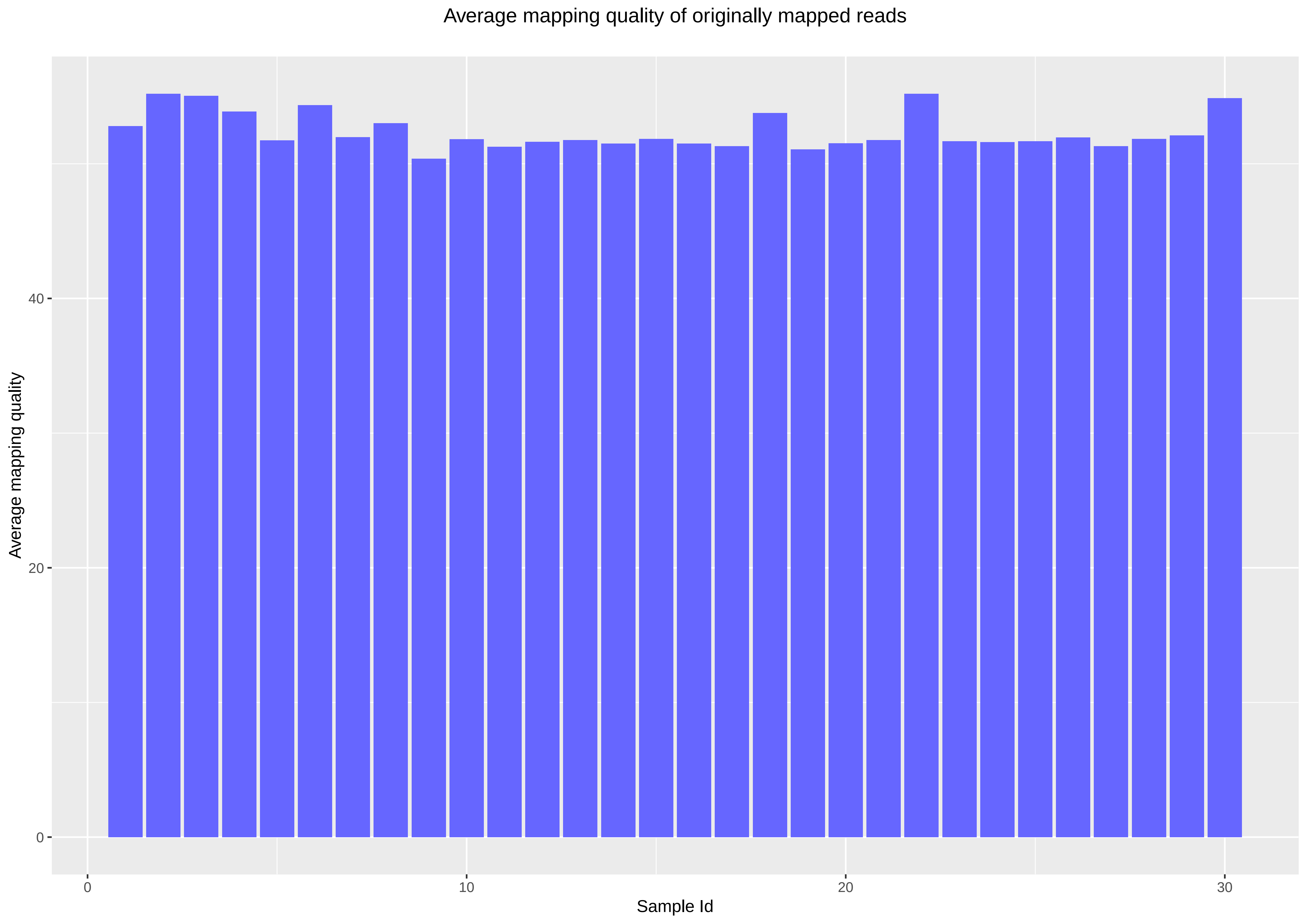
